# Small regulatory RNAs are mediators of the *Streptococcus mutans* SloR regulon

**DOI:** 10.1101/2023.06.02.543485

**Authors:** India Y. Drummond, Alessandra DePaolo, Madeline Krieger, Heather Driscoll, Korin Eckstrom, Grace A. Spatafora

**Author notes:** Address correspondence to Grace Spatafora. Present address: Madeline Krieger, Knight Cancer Institute and Center for Early Cancer Detection, School of Medicine, Oregon Health and Science University, Portland Oregon, USA.

## Abstract

Dental caries is among the most prevalent chronic infectious diseases worldwide. *Streptococcus mutans*, the chief causative agent of caries, uses a 25 kDa manganese dependent SloR protein to coordinate the uptake of essential manganese with the transcription of its virulence attributes. Small non-coding RNAs (sRNAs) can either enhance or repress gene expression and reports in the literature ascribe an emerging role for sRNAs in the environmental stress response. Herein, we identify 18-50 nt sRNAs as mediators of the *S. mutans* SloR and manganese regulons. Specifically, the results of sRNA-seq revealed 56 sRNAs in *S. mutans* that were differentially transcribed in the SloR-proficient UA159 and SloR-deficient GMS584 strains, and 109 sRNAs that were differentially expressed in UA159 cells grown in the presence of low versus high manganese. We describe SmsR1532 and SmsR1785 as SloR- and/or manganese-responsive sRNAs that are processed from large transcripts, and that bind SloR directly in their promoter regions. The predicted targets of these sRNAs include regulators of metal ion transport, growth management via a toxin-antitoxin operon, and oxidative stress tolerance. These findings support a role for sRNAs in coordinating intracellular metal ion homeostasis with virulence gene control in an important oral cariogen.

**IMPORTANCE:** Small regulatory RNAs (sRNAs) are critical mediators of environmental signaling, particularly in bacterial cells under stress, but their role in *Streptococcus mutans* is poorly understood. *S. mutans,* the principal causative agent of dental caries, uses a 25 kDa manganese-dependent protein, called SloR, to coordinate the regulated uptake of essential metal ions with the transcription of its virulence genes. In the present study, we identified and characterize sRNAs that are both SloR- and manganese-responsive. Taken together, this research can elucidate the details of regulatory networks that engage sRNAs in an important oral pathogen, and that can enable the development of an effective anti-caries therapeutic.

## INTRODUCTION

*Streptococcus mutans* is a ubiquitous obligate biofilm-forming bacterium that colonizes the oral cavities of children and adults worldwide (1, 2). It is considered to be the primary etiologic agent of dental caries, owing to its biofilm lifestyle and acidogenic and aciduric properties (2–5). As one of the most common chronic infectious diseases (6, 7), dental caries remains a major public health concern, especially among the socioeconomically-disadvantaged, and it reflects the huge disparities in access to oral and general health care across society (8).

*S. mutan*s is an important member of the diverse healthy plaque microbiome (9–11). However, when homeostatic conditions are disrupted, as they are by a Western diet high in sugar, *S. mutans* can generate lactic acid as a byproduct of sucrose metabolism (2, 4). This acid production drops the pH of the plaque biofilm, thereby favoring the growth of *S. mutans*, which can thrive at a pH as low as 4.0 (12). The ecology of the plaque microbiome therefore shifts towards a state of dysbiosis that favors the demineralization of tooth enamel and the onset of decay (3, 7, 9–11, 13). Taken together with its biofilm lifestyle, the ability of *S. mutans* to both generate acid and survive in its presence enables it to readily transition from a commensal member of the healthy plaque microbiome to a caries-causing pathogen.

The prevalence of *S. mutans* in dental plaque is also due, in part, to its ability to successfully establish and maintain essential intracellular metal ion homeostasis, even as exogenous metal ion availability fluctuates between and during mealtimes. The bacterium maintains homeostasis by tightly modulating metal ion transport across the cell surface. Specifically, *S. mutans* is endowed with at least two metal ion uptake systems, the SloABC Mn^2+^ transport system and the MntH Nramp-type transporter, both of which are upregulated in response to conditions of low Mn^2+^ that prevail between mealtimes (14–16). When exogenous metal ions are plentiful however, such as during a mealtime, *S. mutans* can down-regulate its metal ion-transport machinery to prevent metal ion over-accumulation and the production of toxic oxygen radicals by Fenton chemistry (17–19).

*S. mutans* modulates manganese uptake via its manganese-dependent SloR transcriptional regulator, which also regulates virulence gene transcription across the genome (14, 15, 20–25). SloR is the product of a 654 bp *sloR* gene which is transcribed as part of a polycistronic mRNA via transcriptional readthrough from a promoter located upstream of the *sloABC* operon, as well as independently from its own promoter located in the intergenic region between the *sloC* and *sloR* genes (26, 27). SloR binds to manganese, when it is available, to promote SloR homodimerization and subsequent DNA binding to so-called <u>S</u>loR <u>R</u>ecognition <u>E</u>lements (SREs) (22, 28) in the *sloABC* and *mntH* promoter regions. This SloR-SRE interaction leads to transcriptional repression of the *sloABC* and *mntH* genes (14–16, 27), thereby allowing *S. mutans* to avoid manganese over- accumulation and its associated toxicity.

Small RNAs (sRNAs) are critical regulatory elements that have been a focus of investigation in eukaryotes but remain relatively unexplored in prokaryotes. They are short (∼18- 500 nt), often non-coding transcripts that can function as either enhancers or repressors of gene expression (29–32). Importantly, sRNAs can mediate prompt and energetically low-cost gene regulation post-transcriptionally (33–37). They have been shown to rapidly foster adaptations to environmental stressors such as nutrient limitation and reactive oxygen species in a variety of bacterial pathogens, and they are known regulators of sugar metabolism in *S. mutans* (31, 35, 38, 39). We propose that *S. mutans* sRNAs can coordinate a response to low manganese in the human mouth (i.e., between mealtimes) via the simultaneous upregulation of genes that encode metal ion scavenging proteins and virulence attributes. Such a role for sRNAs can ensure *S. mutans* fitness in the transient environment of the human mouth and elucidate a level of nuanced control for SloR-mediated gene expression that has never before been described.

In the present study, we identified sRNAs that were differentially expressed in the wild-type *S. mutans* UA159 strain compared to its GMS584 SloR-deficient mutant and sRNAs that exhibit differential expression in conditions of low versus high manganese; a subset of these sRNAs were both SloR- and manganese responsive. Two sRNAs, SmsR1532 and SmsR1785, were of particular interest because of their location on the genome. SmsR1532 lies immediately downstream of the *sloABCR* operon, while SmsR1785 maps to a set of genes that encode a toxin-antitoxin system. We focused our attention on these two sRNAs by investigating their putative involvement in manganese-dependent SloR-mediated gene expression and exploring how they contribute to *S. mutans* fitness and virulence potential. Taken collectively, this investigation provides a novel mechanism for SloR-mediated gene regulation and introduces sRNAs and their targets as potential candidates for therapeutics development aimed at alleviating *S. mutans*-induced disease.

## RESULTS

### RNA sequencing supports sRNA transcription that is SloR- and/or manganese-responsive

To identify sRNAs that are differentially expressed in *S. mutans* UA159 and its SloR-deficient GMS584 derivative or in UA159 grown under conditions of low versus high manganese, we generated UA159 and GMS584 sRNA libraries (n=4) and sRNA libraries from UA159 cultures grown in SDM containing 1.7 uM or 75 uM MnSO4 (n=3) using an Illumina TruSeq Small RNA Prep Kit per the manufacturer’s instructions. The total number of reads per library ranged from 2 to 13 million, with up to 98% of transcripts mapping to the *S. mutans* UA159 reference genome and up to 30% mapping to within intergenic regions (IGRs) (Fig. 1, Table S1).

**Figure 1.**
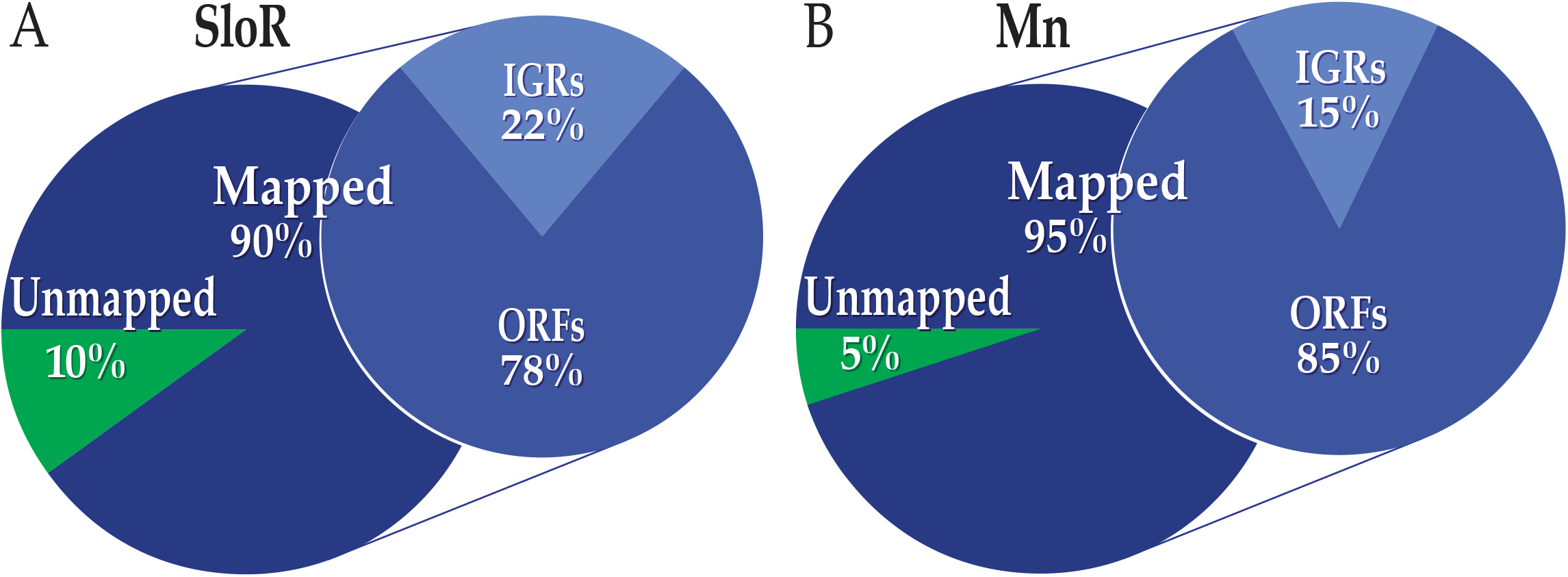
sRNA transcripts derived from RNA-seq. **A)** Shown is the percentage of sRNA transcripts in *S. mutans* UA159 and GMS584 that map to the UA159 reference genome. **B)** Shown is the percentage of sRNA transcripts in *S. mutans* UA159 grown in a medium supplemented with 75 uM (high) versus 1.7 uM (low) manganese that map to the UA159 reference genome. **A and B)** Also shown is the proportion of sRNAs that map to open reading frames (ORFs) or intergenic regions (IGRs) on the *S. mutans* UA159 chromosome.

We identified 56 transcripts 18-50 nt in size in IGRs that were significantly differentially transcribed (p<0.05) in the UA159 vs. GMS584 libraries and 109 transcripts that were significantly differentially transcribed (p<0.05) in the low vs. high manganese libraries. Nine of these transcripts were present in both the SloR and manganese libraries (Fig. 2; Table S4). The reads were then manually filtered to remove sRNAs that mapped to rRNA or tRNA and to select for those sequences with log2 fold change >1.0. From this filtered data we selected 19 sRNAs from the SloR libraries and 10 sRNAs from the manganese libraries that appeared to be independently transcribed according to the bam files (i.e., they had distinct 5’and 3’ ends and their transcripts shared no overlap with other sRNA transcripts (Table S5)). The SloR-modulated sRNAs localized to 14 different IGRs, suggesting that their distribution across the UA159 genome is widespread. Six sRNAs aligned to transcripts that had been previously described in the literature (Table S2-4). Manganese-responsive sRNAs localized to 9 different IGRs across the UA159 genome, with 4 transcripts aligning to transcripts defined previously in the literature (Table S2-4). In the present study, we selected two sRNAs, SmsR1532 and SmsR1785, for further investigation because of their proximity to the *sloABCR* operon and a toxin-antitoxin system, respectively (Table S5).

**Figure 2.**
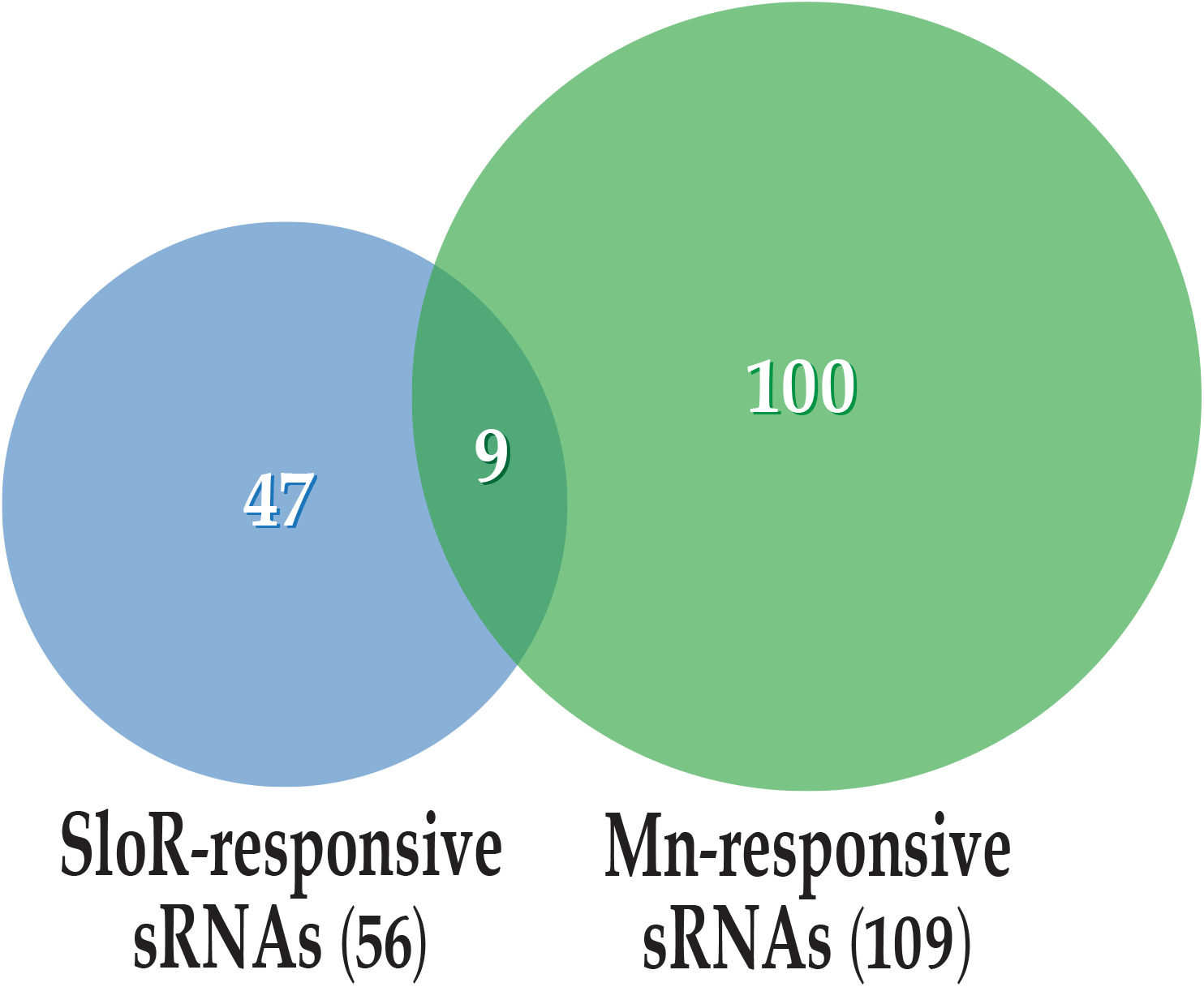
sRNA transcripts can be categorized as SloR- and/or manganese-responsive. We used bioinformatic approaches to reveal 56 sRNAs that are subject to SloR control and 109 sRNAs that are manganese responsive. Among these sRNAs, 9 are positioned at the intersection of SloR and manganese control.

### Organization of SmsR1532 and SmsR1785 on the UA159 chromosome suggests mechanisms for regulatory action

To elucidate the organization of SmsR1532 and SmsR1785 on the *S. mutans* UA159 chromosome, we used both manual and bioinformatic approaches to predict the sequence of the sRNA and its flanking regions. The 5′ and 3′ ends of the predicted sRNAs were determined by junctures in the bam alignment files where read copy number sharply decreased. Since sRNAs are generally not translated and hence do not host the sequencing signals for ribosome or initiator amino-acyl tRNA binding (with the exception of dual-function sRNAs), we used signature sequences of transcription (i.e., promoter elements) and putative SloR recognition elements (SREs) to guide our sRNA characterization.

SmsR1532 is a 30 nt sRNA that localizes to an IGR immediately downstream of the *S. mutans sloR* gene (Fig. 3A). The *sloR* gene is transcribed along with the upstream *sloABC* operon which together encode proteins that transport manganese into the cell and help maintain essential intracellular metal ion homeostasis. The location of this sRNA was of interest because of its downstream proximity to the *sloABCR* genes and thus the potential impact it can have on SloABC- mediated metal ion uptake (40). SmsR1785 is a 66 bp sRNA that maps to an IGR between the *S. mutans* SMU_219 and SMU_220c genes and upstream of the *fst-Sm* gene that encodes a Fst-Sm toxin in *S. mutans* (Fig. 3B) (41). A 56 bp terminator was predicted at the 3′ end of SmsR1532 using ARNold, and a manual search of SmsR1785 and its surrounding sequence revealed a potential 15 bp terminator at its 3′ end (41). mFold predictions support the presence of stable secondary structure within both SmsR1532 and SmsR1785 (Fig. S1).

**Figure 3.**
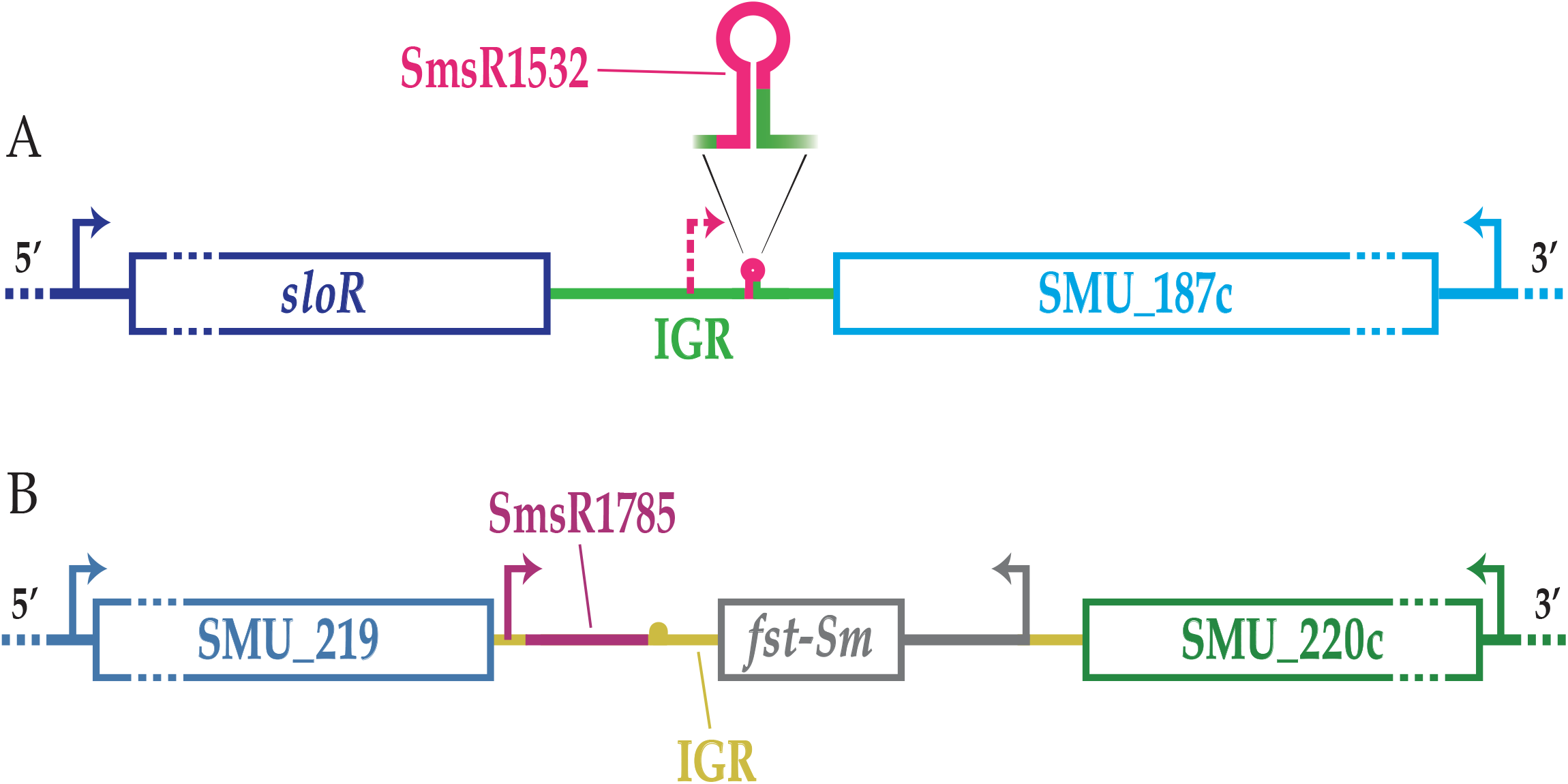
sRNA organization on the *S. mutans* chromosome. Shown are the loci on the *S. mutans* UA159 chromosome where SmsR1532 and SmsR1785 reside. Promoters are represented by the bent arrows and terminator sequences by small loop-out structures. **A)** SmsR1532 is a 3′ UTR-derived sRNA located in the IGR between the *sloR* gene and SMU_187c. The 30 nt transcript (shown in pink) lies within a predicted terminator and is transcribed from a predicted promoter (dashed arrow) in the same direction as *sloR* and antiparallel to SMU_187c. **B)** SmsR1785 is a 3′ UTR-derived antitoxin sRNA that is in the IGR between SMU_219 and SMU_220c and is transcribed in the same direction as SMU_219, but divergent to the *fst-Sm* toxin gene. The predicted terminator begins immediately downstream of the sRNA with a one base pair overlap shared by the sRNA and the terminator (41).

### Coverage plots and qRT-PCR expression profiling experiments confirm sRNA transcription that is SloR- and Mn-responsive

Coverage plots for sRNA libraries generated from *S. mutans* UA159 and GMS584 and UA159 grown in conditions of low and high manganese support the transcription of SmsR1532 and SmsR1785 as 30 nt and 66 nt transcripts, respectively (Fig. 4; Fig. S3). The results of qRT-PCR with a ThermoFisher Custom Taqman Small RNA Assay kit validate the differential transcription of SmsR1532 and SmsR1785 that we noted in the *S. mutans* RNA-seq platform (Fig. 5). That is, transcription of SmsR1532 was de-repressed 5-fold in the SloR-deficient GMS584 strain compared to its SloR-proficient UA159 progenitor (Fig. 5A). In contrast, the expression of SmsR1532 did not change in response to low versus high manganese concentration (Fig. 5B). SmsR1785 expression was 3-fold greater in UA159 compared to GMS584 (Fig. 5C) and nearly 3-fold greater in *S. mutans* UA159 cultures grown in a manganese- replete versus a manganese-deficient medium (Fig. 5D). These results are consistent with a role for sRNAs as small as 30 nt in streptococcal biology.

**Figure 4.**
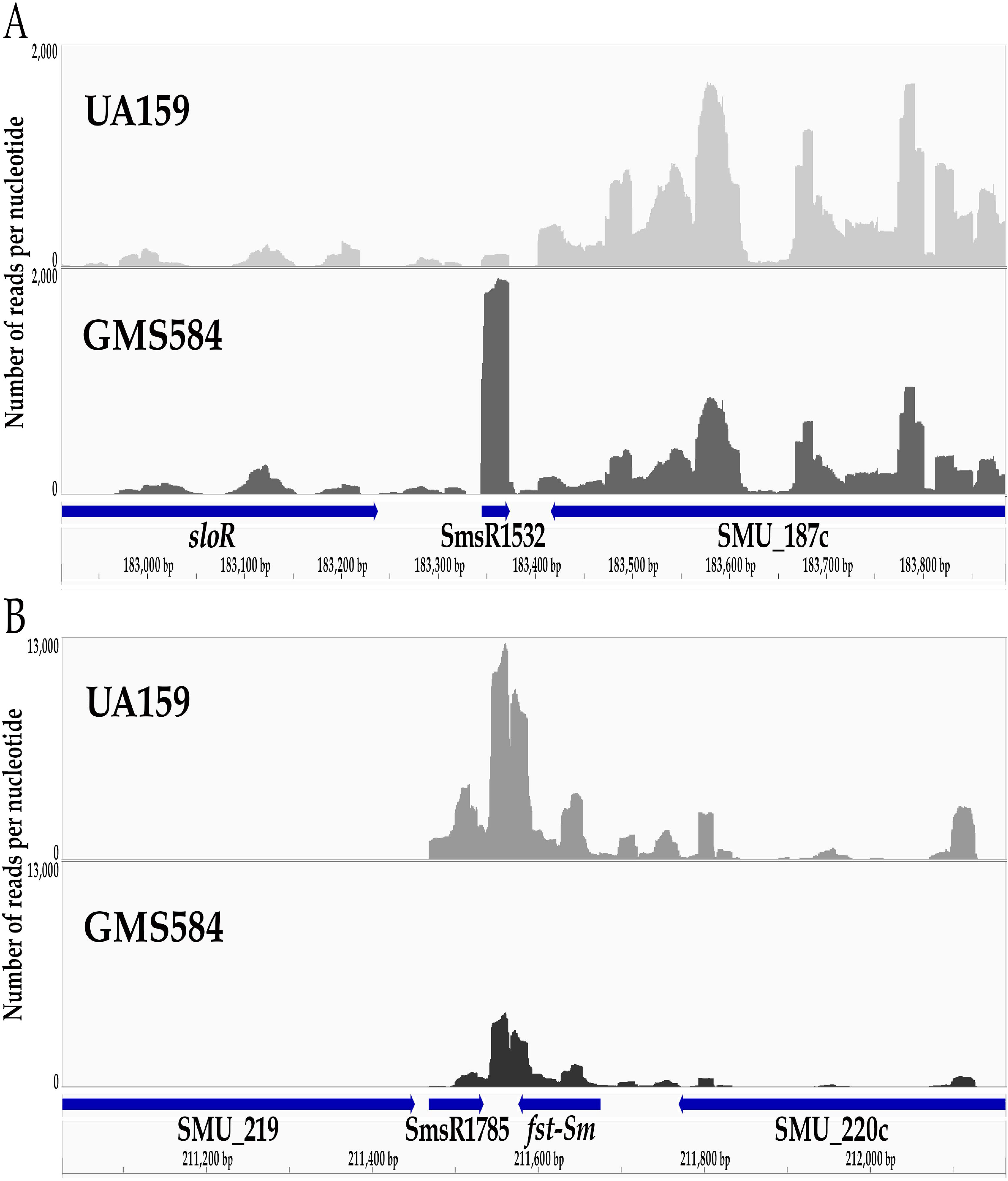
Coverage plots for SmR1532 and SmsR1785 sRNAs in *S. mutans*. Shown are the sRNA-seq reads in the *S. mutans* UA159 wild-type strain and its isogenic SloR-deficient GMS584 mutant. Represented on the y-axes is the number of RNA-seq reads that mapped to each nucleotide. Shown below the coverage plots are the gene and sRNA annotations on the *S. mutans* genome. **A)** SmsR1532 is a 30 nt sRNA that is transcribed from the IGR downstream of the *sloR* gene. **B)** SmsR1785, a 66 nt sRNA, is the previously described srSm antitoxin sRNA in *S. mutans* (41). This sRNA is located downstream and adjacent to the *fst-Sm* toxin gene.

**Figure 5.**
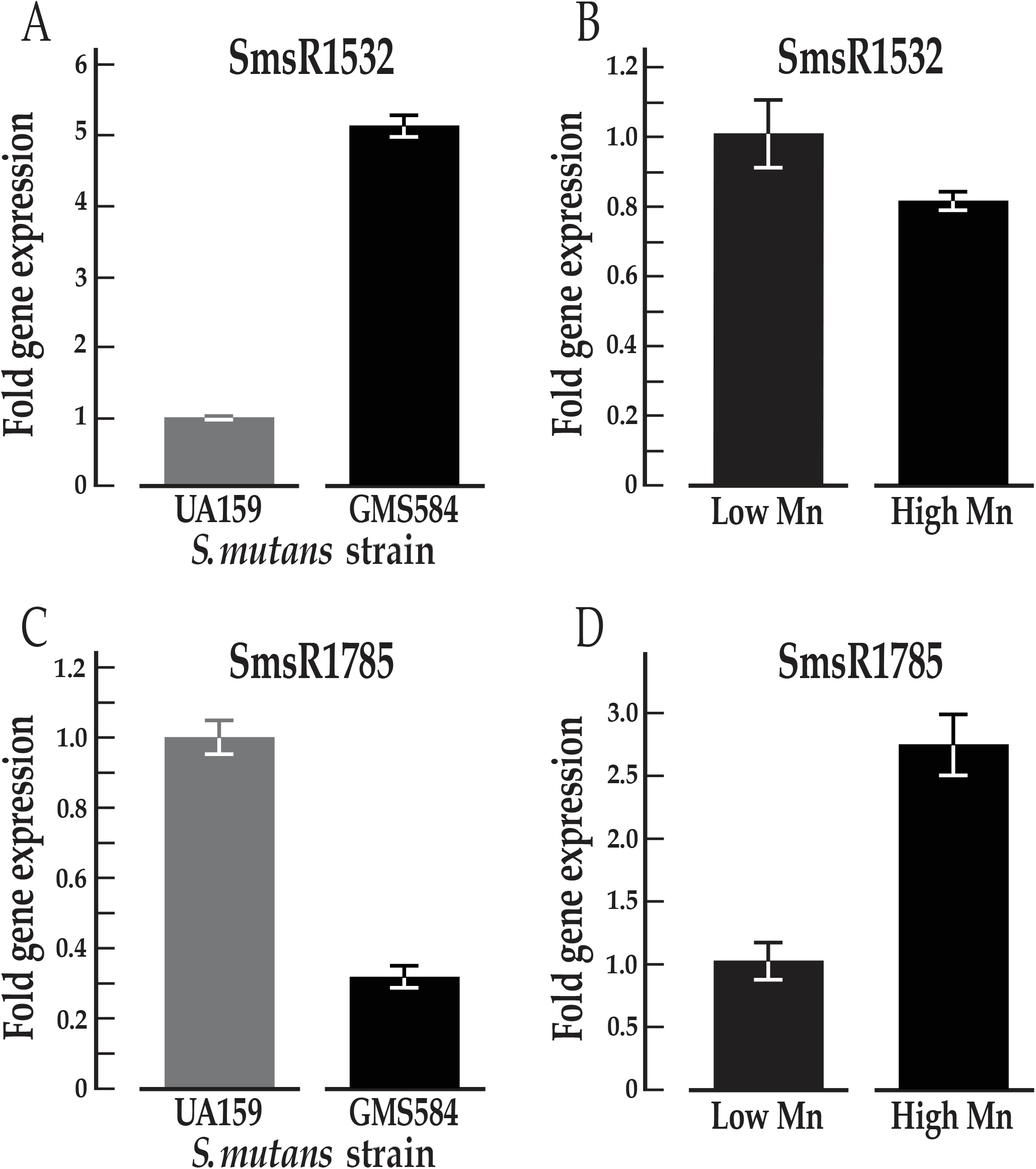
The results of expression profiling experiments validate the output of RNA-seq. Taqman™ qRT-PCR experiments support SloR as a repressor of SmsR1532 and an enhancer of SmsR1785 transcription in *S. mutans*. Parallel experiments support heightened transcription of SmsR1785 at high manganese concentrations. Error bars represent the standard error about the mean. **A)** Expression of *S. mutans* SmsR1532 was upregulated 5-fold in the SloR-deficient GMS584 strain compared to UA159 (n=3, p=1.2x10^-5^ by Student’s *t*-test). **B)** The transcription of SmsR1532 in *S. mutans* UA159 was not manganese-responsive (n=3, p=0.13 by Student’s *t*-test). **C)** Expression of *S. mutans* SmsR1785 was upregulated 3-fold in the SloR-proficient wild-type strain (UA159) compared to GMS584 (n=3, p=4.6x10^-4^ by Student’s *t*-test). **D)** Transcription of *S. mutans* SmsR1785 was nearly 3-fold greater when manganese was present in the culture medium at 75 uM compared to when manganese was limiting (1.7 uM; n=3, p=4.6x10^-3^ by Student’s *t*-test). The qRT-PCR results for UA159 and GMS584 were normalized against the expression of SmsR1491, whereas SmsR847 was used to normalize transcription in UA159 grown in the presence of low versus high manganese; the expression of these sRNAs did not change under the test conditions.

### Comparative genomics and covariance modeling support SloR and sRNA conservation across streptococci as well as unique ancestral origins for SmsR1532 and SmsR1785

*In silico* searches using BLASTN to query a database of 1,053 complete streptococcal genomes (Table S7) revealed that *sloR* is well conserved in the genomes across the streptococci, appearing in 46% of the Streptococcus genomes surveyed and in 70% of the Streptococcus sp. queried (Fig. 6A; Table S12). The *sloR* gene was present in all *S. mutans* strains in the test database. SmsR1532, which we propose could be involved in regulating the adjacent *sloR* and *sloABC* genes, is uniquely present in the representative *S. troglodytae* genome and in 96% of the *S. mutans* strains queried (Fig. 6A; Table S12). Both *sloR* and SmsR1532 were present in 96% of the 25 *S. mutans* strains that we searched (Table S12).

**Figure 6.**
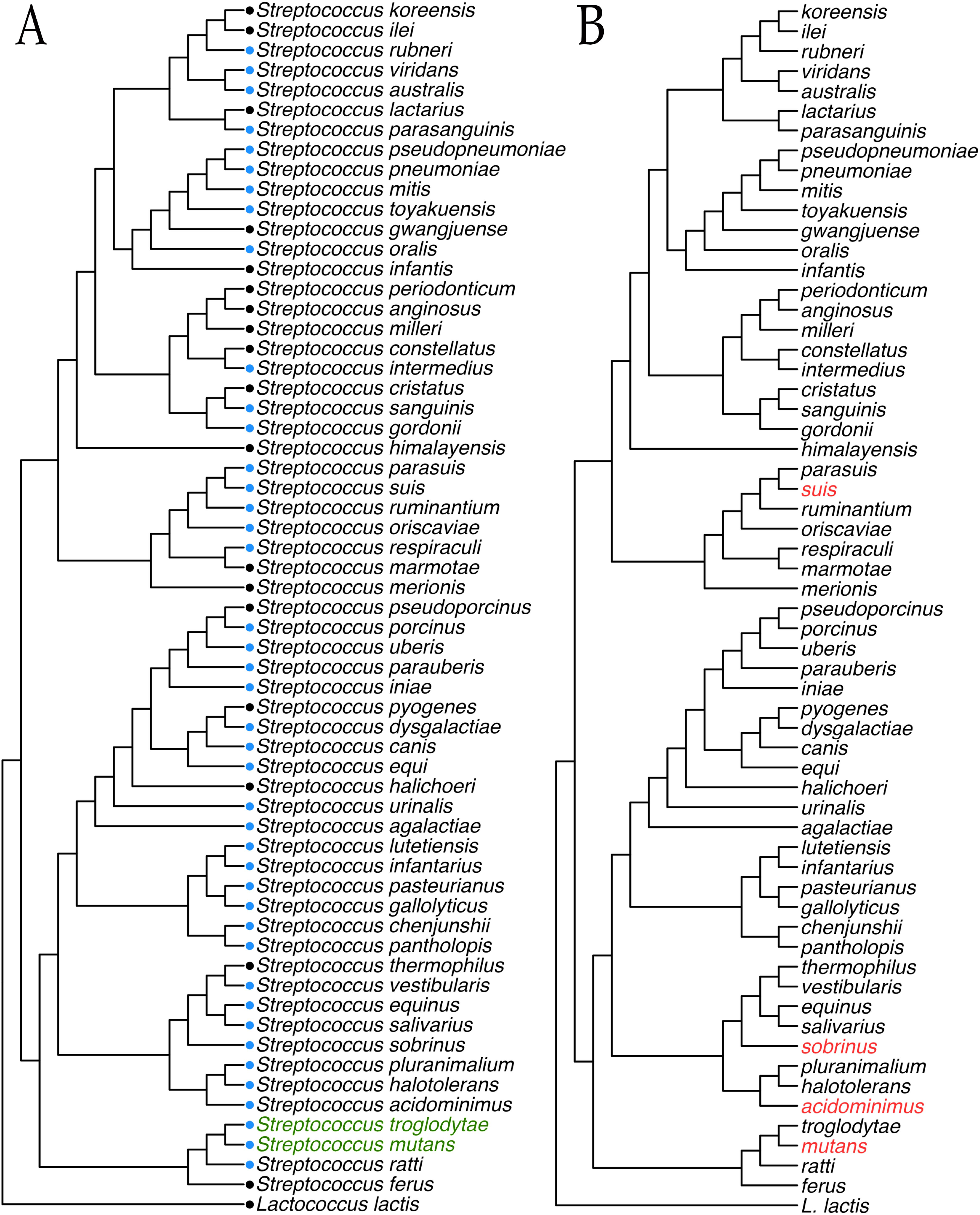
Prevalence of the *sloR* gene and homologs of SmsR1532 and SmsR1785 in streptococci. A cladogram was generated based on single-copy core genes that are present in all 60 representative Streptococcus genomes and rooted in *L. lactis*. **A)** The blue dots indicate the prevalence of the *sloR* gene homologs, and the green text indicates the presence of SmsR1532 homologs as determined by BLASTN. **B)** The red text indicates the presence of SmsR1785 homologs detected through covariance modeling.

Homologs of SmsR1785 were detected in 16% of the *S. mutans* strains in the database, and also in the genomes of *S. suis* and *S. acidimonius* (Fig. 6B; Table S12). All known SmsR1785 homologs locate adjacent to previously annotated toxin-antitoxin system components (Table S11). Interestingly, three copies of the SmsR1785 homolog were present in the genomes of all six *S. sobrinus* strains represented in the database. The location of these sRNAs appears to be very well conserved: one is present in the IGR between a toxin-antitoxin component and a replication-associated recombination protein A gene, another in an IGR between another toxin-antitoxin component and a glucose-6-phosphate isomerase gene, and the third in an IGR between a D-alanine-D-alanine ligase gene and either a ISNCY family transposase or an acyl-CoA thioesterase gene (Table S11).

### Northern hybridization experiments support sRNAs that are resident on large RNA transcripts

To determine whether SmsR1532 and SmsR1785 are transcribed and processed from larger precursor transcripts, we performed northern hybridization experiments with 30 nt and 50 nt complementary probes that span the sRNA transcripts and align at their 5’ ends. The results revealed large hybridizing bands of 106 nt and 73 nt for SmsR1532 and SmsR1785, respectively (Fig. S4). These relatively large RNAs likely represent precursor transcripts that undergo processing to generate smaller mature sRNAs. Processing of the 106 nt RNA to generate a mature 30 nt SmsR1532 product was not visible on the northern blot, suggesting that the blot did not capture transcripts as small as 30 nt and/or that processing intermediates may not exist. In contrast, the SmsR1785 northern blot revealed multiple hybridizing bands ∼73 nt, ∼70 nt, and ∼66 nt in size, consistent with stepwise processing of the 73 nt RNA to yield a 70 nt intermediate and a 66 nt final product. These findings corroborate those in a previous report that describes similar processing of SmsR1785 (41).

### SloR binds directly to the SmsR1532 and SmsR1785 promoter regions

To determine whether SloR modulates sRNA transcription via direct binding, we performed EMSA experiments with the SloR protein and PCR amplicons as target probes. The results of these assays support direct SloR binding to the promoter regions of both SmsR1532 and SmsR1785 (Fig. 7). The target probes in these experiments are shown in Figure 8 and include the sequence of the predicted sRNA and its upstream promoter region, as well as the predicted SRE (Fig. 8). The predicted SRE in the SmsR1532 promoter region directly overlaps the -10 element and the 5′ end of the sRNA. In contrast, the predicted SRE at the SmsR1785 locus is situated promoter-distal, approximately 45 nt upstream of the -35 promoter element. We observed a robust band shift for both SmsR1532 and SmsR1785 when 200 nM SloR was added to the reaction mixture (Fig. 7). The band shift was abrogated when 15 mM EDTA was added, consistent with SloR-DNA binding that is manganese- dependent.

**Figure 7.**
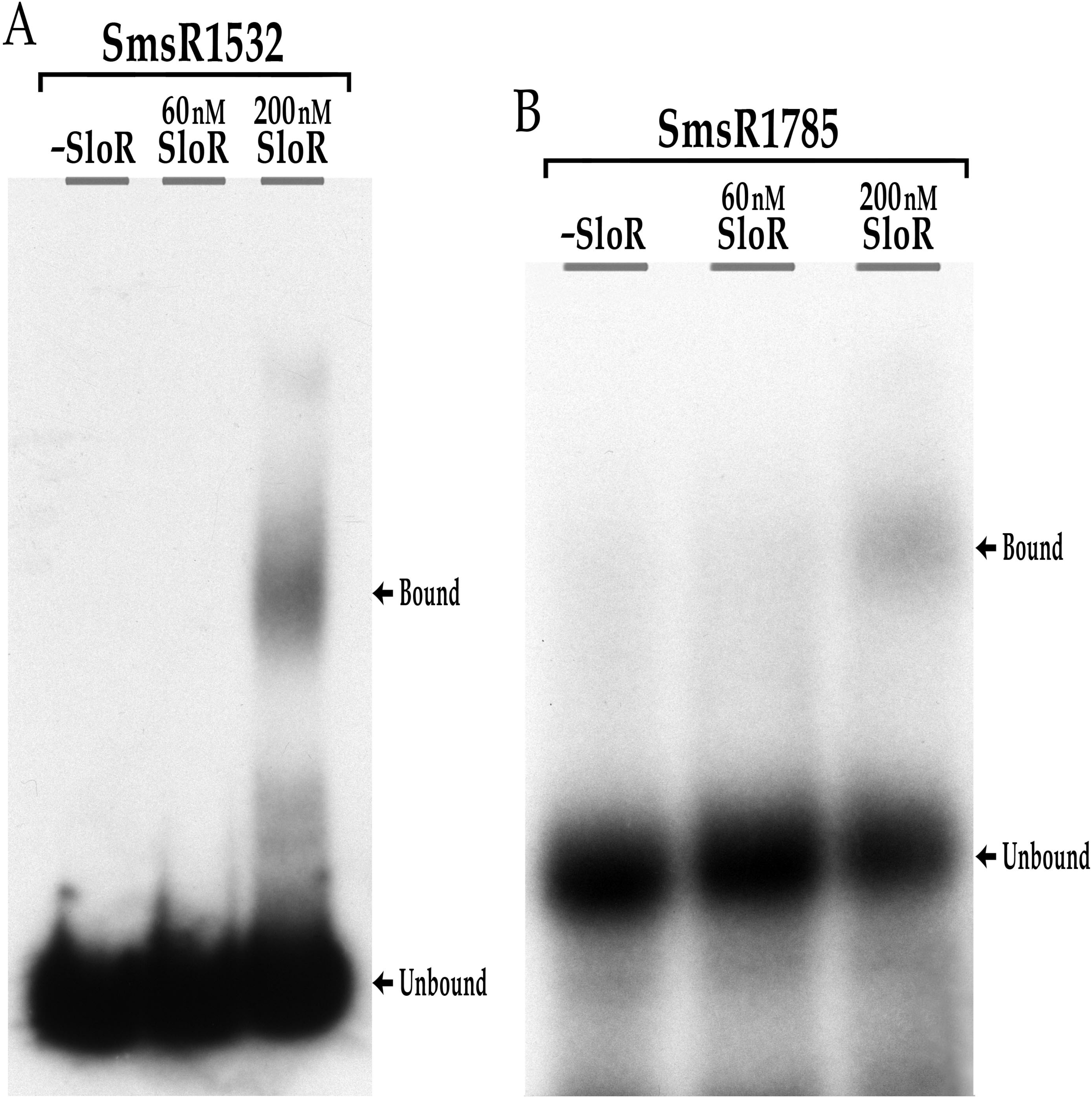
EMSA experiments support a SloR-DNA binding interaction. PCR amplicons containing SmsR1532 **(A)** or SMSR1785 **(B)** along with their upstream promoter regions were end-labeled with gamma 32P and used as target probes. Both probes were shifted in reaction mixtures containing 200 nM SloR on nondenaturing polyacrylamide gels (12%). The addition of EDTA to the SmsR1532 and SmsR1785 reaction mixtures abrogated the band shift (data not shown), confirming SloR-DNA binding that is Mn-dependent. All reaction mixtures were resolved at 300 V for ∼1 hour. Gels were exposed to Kodak XAR film at -80℃ in the presence of an intensifying screen before autoradiography.

**Figure 8.**
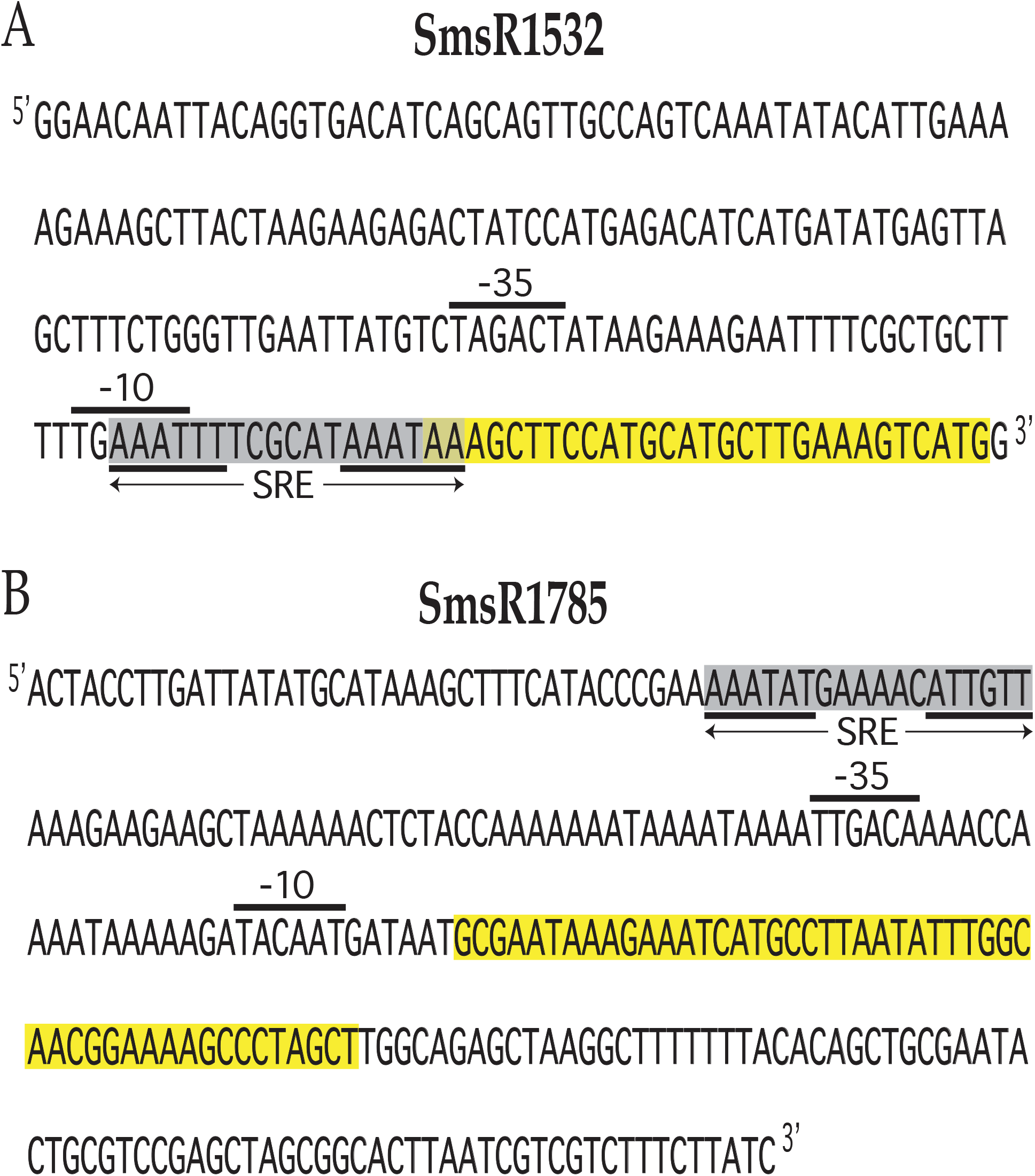
Predicted SloR recognition elements (SREs) map to the SmsR1532 and SmsR1785 promoter regions. (A) A putative SRE with a 6-6-6 binding motif was identified in the SmsR1532 promoter region, sharing overlap with the predicted -10 promoter element. This structural organization is consistent with SloR-SRE binding that blocks promoter access to RNA polymerase, thereby repressing sRNA transcription. **(B)** A putative SRE with a 6-6-6 binding motif was identified distal to the SmsR1785 promoter region, consistent with a SloR-SRE binding interaction that encourages downstream sRNA transcription by RNA polymerase. Shown in **(A)** and **(B)** are the predicted SREs highlighted in gray with the hexameric palindromic repeats underlined. The predicted -10 and -35 promoter elements are indicated upstream of the sRNA sequence, which is highlighted in yellow.

**Figure 9.**
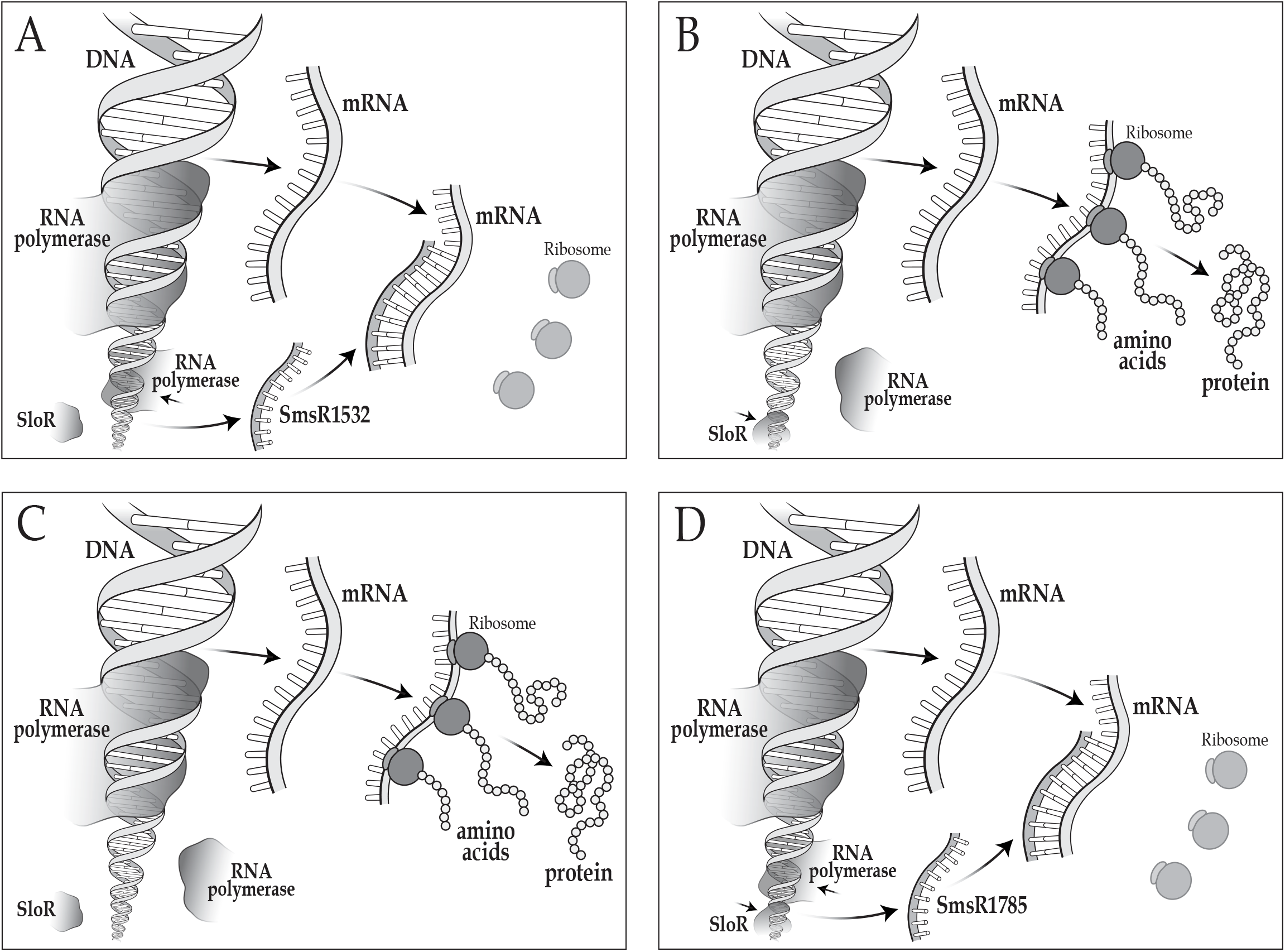
sRNAs are intermediates of the SloR regulon. Shown are scenarios in which SloR may function as an inactive **(A)** or an active **(B)** repressor of SmsR1532 transcription. **A)** During periods of famine, when manganese is limited, SloR cannot readily bind its manganese cofactor and therefore cannot bind DNA. RNA polymerase is free to bind to the DNA at promoter locations that drive transcription of the SmsR1532 sRNA and the mRNA that it targets. The sRNA is then available to bind to its target mRNA via complementary base pairing which can block the ribosome binding site (unlabeled) so that translation cannot occur. **B)** During periods of feast, when manganese is available to bind SloR, the protein can form a SloR-Mn^2+^ complex that binds to the promoter-proximal SRE and blocks RNA polymerase from transcribing SmsR1532. The target gene, however, is transcribed and in the absence of the sRNA, the mRNA is translated. **C** and **D)** Shown are two scenarios in which SloR may act to facilitate SmsR1785 transcription. **C)** When manganese is limiting (e.g., famine), SloR cannot bind to DNA. Consequently, the interaction of RNA polymerase with the SmsR1785 promoter is weakened and transcription of the SmsR1785 is minimal and not readily available to engage with its target mRNA. The mRNA is translated into protein. **D)** When Mn^2+^ is available (e.g., feast), SloR-Mn^2+^ complexes dimerize and bind to a promoter-distal SRE upstream of the SmsR1785 promoter sequence. Subsequent DNA bending may encourage a SloR-RNA polymerase interaction that strengthens the binding affinity between the protein complex with the SmsR1785 promoter. SmsR1785 is transcribed and available to anneal to its mRNA target, thereby preventing its translation.

### Predicted downstream binding targets support a regulatory role for SmsR1532 and SmsR1785 in *S. mutans* fitness and virulence

We used TargetRNA2 and IntaRNA to reveal the predicted mRNA targets of SmsR1532 and SmSR1785 (Table S13-16), and KEGG Mapper to guide our understanding of their relevance to *S. mutans* fitness and virulence (Table 1). Targets of SmsR1532 include annotated genes that mediate carbohydrate transport and metabolism (*deoB*, SMU_1233; *rpiA,* SMU_1234; *msmE*, SMU_878), cell growth/division (*pbpX*, SMU_889), a stringent response (*citE,* SMU_1020; SMU_1217c), oxidative stress tolerance (*spxA2*, SMU_2084c), metal ion homeostasis (SMU_1934c), and exopolysaccharide (EPS) synthesis and biofilm formation (vicR, SMU_1517) (Table S13-14). Although not present in the genome-wide predictions, *sloA*, *sloB*, *sloC*, and *sloR* were recognized as SmsR1532 targets via IntaRNA. Among the most likely targets of SmsR1785 is the mRNA that encodes the *S. mutans* Fst-Sm toxin, which has been shown to hybridize with an srSm antitoxin. Additional predicted targets include annotated transcripts that code for the YknZ (SMU_864) toxin in *S. mutans*, EPS synthesis and biofilm formation (*vicR*, SMU_1517), surface hydrophobicity, (*dltD*, SMU_1688), and contributors to the acid (*livG*, SMU_1666; *atpA*, SMU_1530) and oxidative (*trxA*, SMU_1869) stress response (Table S15-16).

**Table 1.** Predicted gene targets for SmsR1532 and SmsR1785 identified by TargetRNA2 and IntaRNA. Shown are several mRNA targets that were predicted to interact with SmsR1532 and SmsR1785 by TargetRNA2 and IntaRNA freeware. The predicted targets are consistent with a role for these sRNAs in promoting *S. mutans* fitness and virulence potential. A full listing of predicted mRNA targets for SmsR1532 and SmsR1785 can be found in the supplementary material (Tables S13-16).

| sRNA | Predicted targets | Predicted target roles |
| --- | --- | --- |
| SmsR1532 | ABC transporters | *pst* phosphate transport system (*pstC*)<br>sugar transport system (*msmE*)<br>osmoprotectant permease *opuCb*<br>amino acid transport system<br>cobalt transport system<br>putative permease (2) |
|  | *citE* | stringent response |
|  | *spxA2* | Cell envelope homeostasis, oxidative stress |
|  | *vicR* | EPS synthesis and biofilm formation |
|  | *glmZ* | an sRNA-inactivating NTPase |
| SmsR1785 | Fst-Sm | type I toxin-antitoxin system Fst family toxin |
|  | ABC transporters | *pst* phosphate transport system (*pstS*)<br>a branched chain amino acid transporter (*livG*)<br>putative permease<br>Putative efflux pump for Sporulation-Delaying<br>Protein (SDP) toxin<br>ATP binding protein for uncharacterized ABC<br>transport system |
|  | *dltD* | surface hydrophobicity and IPS accumulation |
|  | *trxA* | oxidative stress |
|  | *atpA* | acid stress |
|  | *vicR* | EPS synthesis and biofilm formation |

## DISCUSSION

Small regulatory RNAs can fine-tune gene regulation so that cells can readily adapt to environmental change at low energetic cost (33–36). The oral cavity, arguably the most highly transient environment in the mammalian host, is home to a multitude of bacterial species that must withstand frequent changes in pH, oxygen content, nutrient availability, and essential metal ion concentration during periods of feast and famine (9, 13). *S. mutans* is among the oral microbes that have evolved sophisticated systems to detect and adapt to these environmental fluctuations, many requiring a rapid gene expression response (1, 3–5).

In the present study, we propose that *S. mutans* uses sRNAs to simultaneously regulate metal ion transport and virulence gene transcription. sRNAs have been shown to play an important role in coordinating metal-ion sensing and virulence gene regulation in other bacteria, such as *Staphylococcus aureus*, which uses a manganese dependent RsaC sRNA to ensure a protective oxidative stress response. Under conditions of low manganese, RsaC is transcribed and available to interact with its target *sodA* mRNA, thereby blocking translation of the SodA superoxide dismutase (42). *S. aureus* therefore switches to its SodM superoxide dismutase for oxidative stress tolerance. Herein, we demonstrate a role for *S. mutans* sRNAs in coordinating environmental manganese sensing and virulence gene expression. Specifically, we identified sRNAs that are controlled by the *S. mutans* SloR metalloregulatory protein and/or by manganese concentration, and that target important downstream gene products that contribute to *S. mutans* metal ion uptake and virulence gene expression. This is the first report to highlight such a role for sRNAs in the *S. mutans* manganese dependent SloR regulon.

The results of sRNA-seq revealed 56 sRNAs in *S. mutans* that were differentially expressed in the presence versus absence of SloR, and 109 sRNAs that were differentially expressed at concentrations of low (1.7 uM) versus high (75 uM) manganese; nine sRNAs were responsive to both SloR and manganese. The manganese concentrations used in the present study reflect levels of free Mn^2+^ in saliva that typically range from 1 nM - 4 uM (43, 44), although they can be as high as 36 uM (45). However, manganese in the plaque environment can reach concentrations as high as 6.2 mM in some cases (46). Adjusting the manganese test conditions to include a broader range of concentrations could elucidate additional novel sRNAs that are SloR- and/or manganese- responsive.

Further sorting of these data revealed 19 SloR-responsive and 10 manganese-responsive sRNAs, two of which were subject to both SloR and manganese control. We were not surprised to observe some sRNAs that were responsive to manganese but not to SloR, since SloR is only one of a multitude of transcription factors in *S. mutans* that is impacted by manganese (19). Given that SloR is a manganese-dependent regulator, however, one might expect all SloR-dependent sRNAs to be Mn-responsive. In fact, not all SloR-responsive sRNAs that we identified were Mn-dependent, a finding that is consistent with reports of other metal ions that can serve as SloR cofactors, such as iron (47). In the present study, we focused our attention on two sRNAs that are SloR and/or manganese-responsive in *S. mutans*, SmsR1532 and SmsR1785.

SmsR1532 is a 3′ UTR-derived sRNA that is located downstream of the *sloR* open reading frame on the *S. mutans* UA159 chromosome. Reports in the literature describe Mn^2+^ transport across the *S. mutans* cell envelope that is modulated by SloR and mediated by the SloB permease which is encoded as part of the *sloABC* operon (14, 15). Interestingly, these genes are located adjacent to and immediately upstream of *sloR* and are transcribed as a polycistron that includes *sloR* as a product of transcriptional readthrough (26, 27). The proximity of SmsR1532 to the *sloABCR* operon is consistent with its putative involvement in *sloABCR* regulation, as sRNAs can readily regulate adjacent genes in *trans* via their seed region. Similar *trans*-acting sRNAs have been described in *Vibrio cholerae* where sRNAs OppZ and CarZ are transcribed from within the 3′ UTRs of their adjacent *oppABCDF* and *carAB* operons, respectively; once the sRNAs are processed and released from their precursor transcripts, they bind to the *oppABCDF* or *carAB* target mRNAs and inhibit their translation (48).

SmsR1785 is a 3′ UTR-derived sRNA that maps to a toxin-antitoxin (TA) locus on the *S. mutans* UA159 chromosome (41). TA systems in bacteria are known to target essential cellular functions like DNA replication, RNA stability, and cell envelope integrity to facilitate microbial fratricide, thereby heightening competitive fitness (49). Adding a TA system to the SloR regulon can broaden our understanding of the SloR metalloregulatory protein and its impact on *S. mutans* virulence gene regulation.

The results of pangenome analysis support SloR as a well-conserved protein across *Streptococcus* sp. In contrast, SmsR1532 appears to have emerged only within the distinct clade containing *S. mutans* and *S. troglodytae*, the latter being a predominant, strongly-adherent, bacitracin-resistant member of the mutans streptococci in the oral cavities of chimpanzees (50). Owing to limitations in the BLASTN algorithm, it is possible that additional Streptococcus species that harbor SmsR1532 may have been overlooked. With that said, SmsR1532 is present in 96% of *S. mutans* strain isolates in the database, all of which derive from a common ancestor.

There is no clear pattern of inheritance within the four *Streptococcus* sp. in which SmsR1785 is present, suggesting acquisition of this sRNA via horizontal gene transfer (HGT). HGT is a common method for exchanging toxin-antitoxin genes across bacteria (51). The near absence of SmsR1785 homologs from *S. mutans* isolates (16%) again supports SmsR1785 acquisition via HGT, indicating that SmsR1785 is not a core component of the *S. mutans* genome. Interestingly, reports in the literature position srSm within a transposon-related island on the *S. mutans* UA159 chromosome (41), lending further support to the HGT hypothesis. The HGT hypothesis is further supported by the location of SmsR1785 in several *S. sobrinus* strains where it is adjacent to an ISNCY family transposase. *S. sobrinus* is unique in that it harbors three copies of SmsR1785. Future studies can elucidate the impact of these redundant sRNAs and whether they speak to the HGT of SmsR1785 within/across the streptococci.

sRNAs that localize to 3′ UTRs can be of two varieties: type I, which is transcribed from an independent proximal promoter, or type II, which derives from the co-transcription with and processing of a larger polycistronic mRNA (40). The results of northern hybridization experiments did not reveal evidence of the 4 kb polycistronic mRNA that would be expected if SmsR1532 were the product of *sloABCR* mRNA processing. Instead, a 106 nt transcript hybridized with the 30 nt SmsR1532 probe, a result that supports sRNA transcription that is driven from an independent promoter. We propose that subsequent processing of the 106 nt SmsR1532 RNA culminates in the 30 nt product that we observed in the sRNA-seq libraries. Other reports in the literature describe similar two-step processing for bacterial sRNAs (36, 40, 52–55). For instance, a highly conserved type I sRNA, called ArcZ, in the Enterobacteriaceae is transcribed as a 120 nt RNA from an independent promoter that resides within the 3′ end of the upstream *arcB* gene (52). This transcript is subsequently processed into a 50 nt sRNA that then goes on to modulate genes that contribute to serine transport and the oxidative stress response.

The results of sRNA-seq support alignment of the 5′ end of SmsR1785 with the transcription start site (TSS) of the previously described antitoxin sRNA (srSm) that was identified by 5′RACE (41). The predicted -35 promoter element of SmsR1785 is located within the coding sequence of the upstream SMU_219 open reading frame, unlike the predicted -10 element which is located immediately downstream of SMU_219. Furthermore, the results support the transcription of SmsR1785 as a 66 nt mature sRNA, as was observed on northern blots. Larger 73 nt and 70 nt precursor transcripts were also visible on these blots, consistent with sequential processing of these RNAs to generate the final 66 nt sRNA. These transcript sizes corroborate those reported previously in the literature (41) and together support the shared identities of SmsR1785 and srSm.

We noted expression of SmsR1532 that was de-repressed in the SloR-deficient GMS584 mutant strain compared to its SloR-proficient UA159 progenitor, indicating that SloR is a repressor of SmsR1532 transcription. SmsR1532 repression aligns with the location of the predicted SRE in the SmsR1532 promoter region, which shares overlap with the predicted -10 promoter element; binding to this SRE would block access of RNA polymerase to the sRNA promoter, thereby repressing SmsR1532 transcription. Indeed, we see this expression pattern for other promoter proximal SREs in the *S. mutans* genome. For instance, the *sloABC* and *mntH* genes, which are SloR-repressed, likewise harbor SREs that share overlap with their respective promoter elements. The results of EMSA support direct SloR binding to the SmsR1532 promoter, which includes a predicted SRE with a palindromic hexameric repeat sequence.

In contrast to SmsR1532 expression, we observed SmsR1785 transcription that was upregulated in the SloR-proficient UA159 strain compared to the GMS584 SloR-deficient strain, suggesting that the presence of SloR encourages the expression of this sRNA. This expression is consistent with the location of the predicted SRE in the SmsR1785 promoter region which is promoter-distal, localizing 95 bp upstream of the SmsR1785 transcription start site TSS. The correlation between promoter distal SREs and genes that are upregulated by SloR is supported by reports in the literature. For example, transcription of the *S. mutans bta* gene is facilitated by SloR binding to a promoter-distal SRE located 167 bp upstream of the *bta* TSS (21). It is possible that SloR-SRE binding at the SmsR1785 and *bta* loci encourages DNA bending to facilitate gene transcription. The results of EMSAs support direct SloR binding to target probes that include the SmsR1785 promoter region and its promoter distal SRE. SmsR1785 expression was also upregulated in *S. mutans* cells grown in the presence of high manganese, an expression pattern that parallels the heightened expression we observed for SmsR1785 in the SloR-proficient UA159 strain. Taken together, we propose that SloR-Mn^2+^ complexes are favored under these test conditions which, in turn, encourage SloR homodimerization, DNA binding and active SmsR1785 transcription.

*In silico* analysis of SmsR1532 and SmsR1785 with Target2RNA and IntaRNA freeware revealed the formation of stable intramolecular hairpin structures (ΔG = -2.40 kcal/mol and ΔG = -17.90 kcal/mol, respectively). However, the positioning of these stem-loop structures in relation to the different mRNA-specific proposed seed regions on the sRNAs does not lead to intuitive sRNA-target interactions. That is, the seed regions of both SmsR1532 and SmsR1785 occur within the stem-loop structure and may therefore be inaccessible for target binding. With that said, the ΔGs that are associated with each of the sRNA-target mRNA pairings are sufficiently more negative than the ΔG that is associated with the intramolecular sRNA secondary structures, thereby favoring the interaction of each respective sRNA with its downstream mRNA target. We propose that the hairpin structure inherent in these sRNAs may therefore serve as a stabilizing factor when the sRNA is in its unbound configuration, and that it functions as a “threshold-setter” for sRNA-mRNA binding when presented with an appropriate binding target. That is, the only sRNA-mRNA target interactions that can occur are those with a ΔG that is more negative than that of the intrinsic sRNA hairpin stabilizer. We propose that these thermodynamics could ensure the melting of the sRNA hairpin and subsequent binding specificity of the sRNA to its intended target.

We went on to apply the IntaRNA and TargetRNA platforms to predict the downstream targets of SmsR1532 and SmsR1785. Among the predicted targets of SmsR1532 are the *sloABCR* genes, which belong to the *S. mutans* SloR regulon. Sequence queries in ARNold revealed a multitude of locations that span the *sloABCR* polycistron where SmR1532 can bind in *trans* to repress translation of the *sloABCR*-encoded metal ion uptake machinery. We propose that these SmsR1532 and *sloABCR* interactions could serve to fine-tune regulation of the *sloABCR* operon beyond the regulation that is dictated by SloR alone. The *S. mutans* Fst-Sm toxin is among the targets predicted for SmsR1785. The location of the seed region in this sRNA, according to IntaRNA, is the site that is known to bind Fst-Sm and prevent Fst-Sm-induced toxicity (41).

Other downstream targets of SmsR1532 include *vicR* and *spxA2*, while targets of SmsR1785 include *fst-sm*, *vicR*, *dltD*, and *trxA*. The *vicR* target of SmsR1532 and SmsR1785 is involved in EPS production (56–58), while the *spxA2* target of SmsR1532 encodes a major regulator of the cell envelope stress response and, to a lesser extent, the oxidative stress response (59–61). The *dltD* target of SmsR1785 is involved in lipoteichoic acid metabolism and plays a role in cell surface hydrophobicity (62, 63) and ultimately the cariogenic potential of *S. mutans* (64). In contrast, the *trxA* target of SmsR1785, is a *S. mutans* thioredoxin and a key player in the oxidative stress response (65). Taken together, these predictions support *S. mutans* sRNAs that, in response to manganese limitation, modulate mRNAs that contribute to *S. mutans* fitness and virulence (e.g., metal ion homeostasis, mutacin production, EPS synthesis, cell envelope stress response, surface hydrophobicity, and oxidative stress tolerance).

## Conclusion

The results of the present study make it increasingly clear that sRNAs play a paramount role in bacterial gene regulation. This study is unique however, in that it links virulence gene control in the oral pathogen, *S. mutans* to environmental cues in the mouth such as metal ion sensing. Specifically, we propose that manganese limitation during extended periods of nutrient deprivation in the oral cavity (i.e., between mealtimes) signals *S. mutans* to upregulate its virulence attributes as it scavenges for essential micronutrients via its metal ion transport machinery. This coordinated control can foster *S. mutans* fitness in the highly transient conditions of the human mouth. The mechanism(s) by which sRNAs fine-tune virulence to optimize *S. mutans* survival and persistence in the mouth, however, will require further investigation. An improved understanding of how *S. mutans* engages sRNAs in virulence gene transcription can reveal novel targets for therapeutics development aimed at alleviating or preventing *S. mutans*-induced caries.

## MATERIALS AND METHODS

### Bacterial strains, growth conditions, and bacteriological media used in the present study

The bacterial strains used in this study included the *S. mutans* wild-type UA159 strain (ATCC 700610) and the UA159-derived, SloR-deficient GMS584 strain (15). For RNA and DNA isolation, *S. mutans* was grown at 37℃ and 5% CO2 with selective pressure when necessary. For expression profiling experiments, the UA159 and GMS584 strains were grown as standing cultures in 15-mL Falcon tubes (Thermo Fisher Scientific) containing Brain Heart Infusion (BHI) media, or in a semi-defined medium (SDM) supplemented with low (1.7uM) or high (75uM) concentrations of MnSO4•H2O (Acros Organics). *S. mutans* GMS584 cultures were grown in the presence of erythromycin (10 ug/mL).

### Primers and oligonucleotides used in the present study

Shown in Table 2 are the primers and oligonucleotide target probes that were used in EMSA and expression profiling experiments.

**Table 2.**
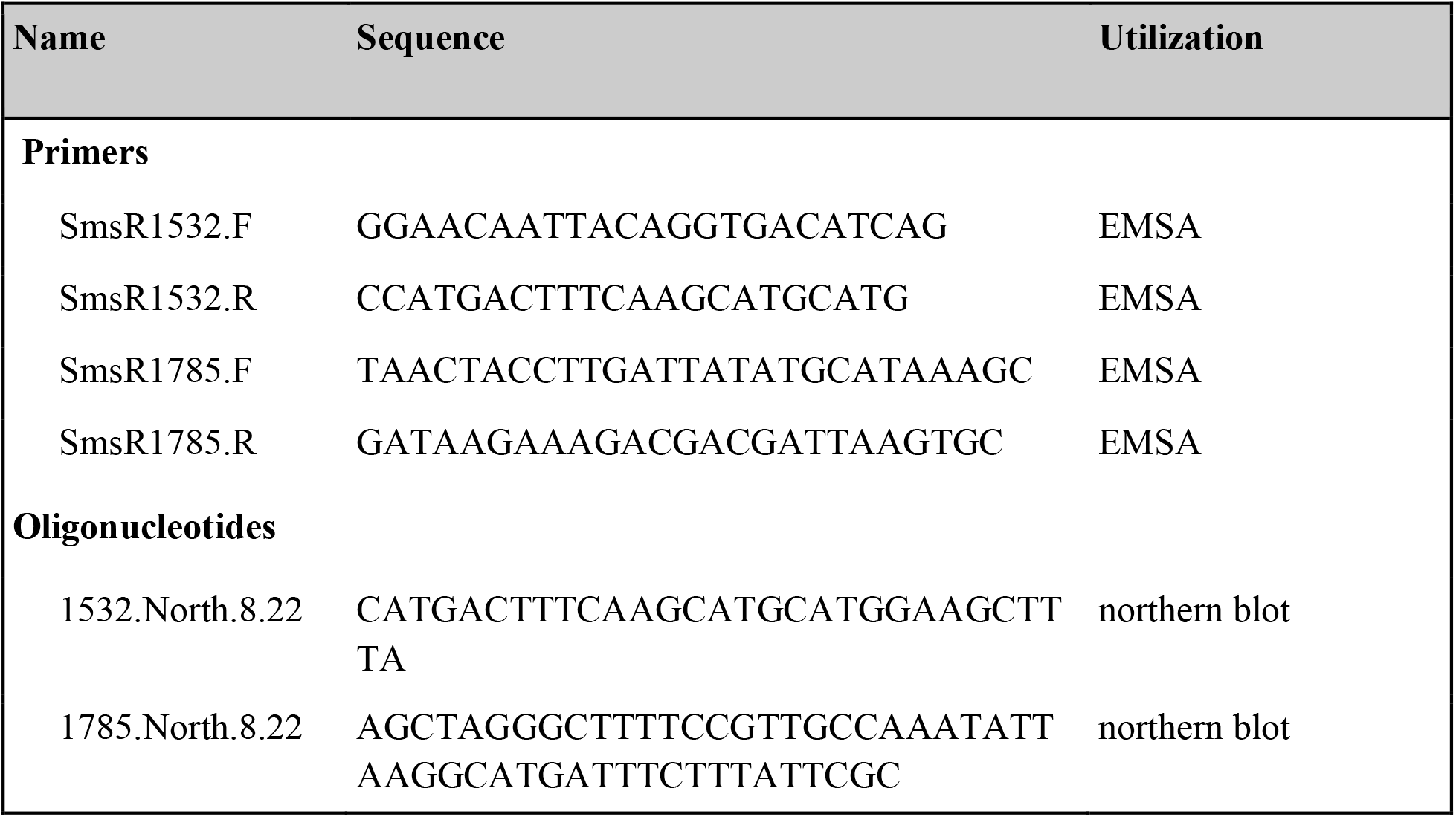
Primer and oligonucleotides used in this study.

### RNA isolation

Total RNA was isolated from *S. mutans* UA159 and GMS584 strains according to a modified protocol of Crepps et al. (23). Briefly, cells from an overnight culture of *S. mutans* were grown to mid-logarithmic phase (OD600nm = 0.4-0.6), pelleted by centrifugation, and resuspended in RNA Protect (Qiagen). The cells were then mechanically lysed and their RNA purified and treated with DNase I using an RNeasy kit according to the manufacturer’s instructions (Qiagen). RNA concentrations were assessed on a NanoDrop Lite spectrophotometer (Thermo Fisher Scientific) and integrity was determined with a Bioanalyzer 2100 (Agilent). All RNA samples were aliquoted and stored at -80℃.

### RNA library construction and sequencing

Four independent, stranded sRNA libraries were generated from *S. mutans* UA159 and GMS584 mid-logarithmic phase cultures using the TruSeq Small RNA Library Prep Kit (Illumina) according to the manufacturer’s instructions. In parallel, three sRNA libraries were prepared for *S. mutans* UA159 cells grown at low and high manganese concentrations as described above. Total RNA was ligated to a 3′ adapter (5′- GUUCAGAGUUCUACAGUCCGACGAUC-3′) and a 5′ adapter (5′-UGGAAUUCUCGGGUGCCAAGG-3′) as described in the TruSeq kit. The adapter-ligated RNA was reverse transcribed using primers provided in the Illumina kit and SuperScript II Reverse Transcriptase (Invitrogen). cDNA was amplified and indexed using the conditions provided in the Illumina protocol; protocol optimization revealed that 18 amplification cycles generated sufficient cDNA concentrations for sequencing without compromising the signal of low abundance sRNAs. Quality of the amplified cDNA was assessed on an Agilent Bioanalyzer 2100. Total cDNA was resolved on a 6% TBE polyacrylamide gel and sRNAs ∼18-50 nt in length were selected by gel extraction. The cDNAs were further purified as described in the Illumina protocol and concentrated in a vacufuge (Eppendorf) at 42℃ for 10-15 minutes. The dried cDNA was resuspended in 10 uL Tris-HCl, pH 8.5, quantified on a Qubit fluorometer (Thermo Fisher Scientific), and quality assessed on an Agilent Bioanalyzer 2100. RNA sequencing was performed at the University of Vermont on an Illumina MiniSeq platform to achieve 7 million paired-end reads 150 bp in length or on a HiSeq 1500 flow cell to achieve 140 million single-end reads 75 bp in length (Table S1)

### A literature-based approach revealed putative sRNAs in *S. mutan*s

A citation database of the peer-reviewed literature called Scopus was used to search for published articles that predict small regulatory RNAs in streptococcal species and in *Staphylococcus aureus* via *in silico* or RNA sequencing (RNA-seq) approaches. Three Scopus searches were performed using the terms “sRNA,” “small regulatory RNA,” and “small RNA,” as well as either “*Streptococcus mutans*,” “*Streptococcus*,” or “*Staphylococcus aureus.*” Available sequences were compiled into a sRNA_MASTER.fa fasta file (Table S2), along with the sRNA name, the bacterial strain, whether the sRNA is located on the chromosome or a plasmid, the transcription start and stop coordinates (if available), and the source study. For those studies that reported sRNA coordinates but not sequences, a perl script was prepared and used to extract the sequences from their reference genomes. The table was then sent to the nearby University of Vermont to determine whether any sRNAs from the published literature were direct matches with the sRNAs identified in the present study by RNA-seq and BLAST. All reports of *S. mutans* sRNAs published as of August 2022 are presented in Table S3.

### Identification of the RNA-seq output for sRNAs in *S. mutans*

The workflow that was used to generate a list of candidate sRNAs is presented in Fig. S1. Briefly, Trim Galore! was used in its mode for sRNA-seq reads to remove low quality reads, reads <18 bp, and adapters. The cleaned reads were mapped to the UA159 reference genome (RefSeq <u>NC_004350.2</u>) using bowtie2 with the following parameters for mapping short reads for the SloR libraries: -n 1 -1 10 -m 100 -k 1. Default parameters were used for the manganese libraries. samtools was used to sort and index the alignment files. Next, sRNA-Detect (66) was used to remove transcripts that map to known coding sequences in the *S. mutans* UA159 genome and to align the remaining transcripts across libraries. These sRNAs were then filtered to select for reads that were present in 6/8 UA159/GMS584 libraries or 4/6 low/high manganese libraries, had >100 reads, and were <50 bp in length. samtools was used to filter the sam files based on the coordinates of intergenic reads such that only transcripts within intergenic regions remained. This filtering step helped to further remove reads that mapped to tRNA, rRNA, and CDS. The alignment files were converted to fastq files using either bam2fastq or Picard sam2fastq, and then served as the input files for DEUS (67). In DEUS, CD-HIT was used to cluster together sRNAs (agnostic of reference genome) that may have derived from the same transcript but degraded to different degrees. For the SloR libraries, a BLASTN analysis was performed to see if any of the clustered sequences aligned with sRNA sequences described in the literature. Differential expression of sRNAs in UA159 versus GMS584 was subsequently assessed. For the manganese libraries, DEUS was first used to assess differentially expressed sRNAs in low and high manganese concentrations. A BLASTN search was then used to determine whether any of these sRNAs aligned with sRNA sequences described in the literature.

Upon manual inspection, we noted redundant transcripts within the data set. We used CD-HIT-EST v4.8.1 to collapse these redundant transcripts (Table S4). CD-HIT-EST was run using the default parameters implemented in DEUS with the exception of the length difference cutoff, which was set to 0.4 to allow for the clustering of shorter sRNAs with longer cluster representatives. The former needed to be at least 40% the length of the latter. sRNA clusters resulting from CD-HIT-EST were used to find the intersection of differentially expressed sRNAs in the datasets visualized as a Venn diagram.

Manual processing promoted the selection of transcripts 18-50 nt in length with a log2 fold change greater than ±1.0 and a p-value <0.05. These sRNAs were mapped to the UA159 genome in Benchling to establish their coordinates, confirm their location in IGRs, and identify adjacent genes as well as general genome location. The genome viewer Tablet (68) was used to view the bam files and determine whether reads represented independent transcripts. sRNAs that were clearly defined in the bam files, >25 nt, and consistently expressed across libraries for a particular strain were selected for further investigation (Table S5).

### sRNA characterization *in silico*

We used mFold web server version 3.6 (V3.6) (69–71) with default parameters to predict the secondary structure of sRNA sequences revealed by sRNA-seq (Fig. S2). The presence of -10 and -35 promoter elements was assessed using a manual scan of the region upstream of the sRNAs. ARNold Rho-independent termination prediction freeware (72–75) was used to identify putative terminator sequences.

### Coverage plots

Bam files were visualized using the Integrative Genomics Viewer (version 2.14.0) using a track height of 400. Data ranges of 2,000 and 13,000 reads per nucleotide were used for SmsR1532 and SmsR1785 expression in the SloR libraries and 45 and 100 reads per nucleotide for SmsR1532 and SmsR1785 expression in the manganese libraries.

### Phylogenetic tree construction

Pangenome analysis was performed using the Anvi’o workflow (76, 77) on 60 complete genomes representing unique Streptococcus species, plus a *Lactococcus lactis* genome that served as an outgroup (Table S6). This pangenome was used to identify 318 single-copy core genes present in all genomes, which were then used to construct a phylogenetic tree using FastTree (78) within the Anvi’o workflow. The tree was rooted in *L. lactis* and visualized in R using the ggtree (version 3.21) and treeio (version 1.18.1) packages.

### BLAST searches

A database containing 1,053 complete Streptococcus genomes downloaded from NCBI (Table S7) was compiled and used for all subsequent BLASTN and covariance modeling searches. The *sloR* sequence from *S. mutans* UA159 was used as a query with default BLASTN parameters, and the results were annotated with an in-house python script (Table S8). All BLASTN hits with previous annotations to a *sloR* gene homolog were retained, and annotated hits were filtered based on e-value (<.05) (Table S9).

Because of its small size (30 nt) and lack of divergent sequences detected in the Streptococcus database, SmsR1532 was not a good candidate for covariance modeling. Instead, homologous sequences were detected via a BLASTN search for small transcripts (word size 7) (Table S8). All hits with 100% sequence identity and >96% coverage were retained (Table S10).

### Covariance modeling

Covariance modeling was performed for SmsR1785 as previously described (37, 79, 80). Briefly, the 66 nt SmsR1785 sequence (41) (Table S8) was used as a BLASTN query to search a database of 1,053 complete Streptococcus genomes. Five unique sequences with 72-91% percent identity and 51-100% coverage of the query were selected and used along with the SmsR1785 UA159 sequence to construct the covariance model. LocARNA (81) was used to align sequences, and the infernal suite of tools (82) (version 1.1.4) was used to construct and calibrate the model (cmbuild and cmcalibrate, respectively). Next, a covariance model search (cmsearch) was performed with the database of 1,053 genomes to reveal new hits with unique sequences and an e-value of <1e−5. These hits were added to the model, which was then recalibrated and used to search the database. This iterative process was repeated until no new hits were detected. The final results were annotated with an in-house python script (Table S11).

### TaqMan Small RNA Assays

Custom Taqman™ Small RNA Assays (Applied Biosystems) were used to both validate and characterize the expression of SmsR1532 and SmsR1785; SmsR1491 and SmsR847 were used as endogenous controls since their expression did not change under the experimental test conditions (i.e., in UA159 versus GMS584 and in UA159 grown in high versus low Mn^2+^, respectively). Small RNA-specific assays were designed using the Custom TaqMan™ Small RNA Assay Design page and, upon company approval, ordered from Applied Biosystems. Two-step RT-PCR was performed according to the manufacturer’s instructions in each of three independent experiments. Briefly, total RNA was isolated from both *S. mutans* UA159 and GMS584 as described above. sRNA-specific reverse transcription was performed with 10 ng of total RNA using the Taqman MicroRNA Reverse Transcription kit and an sRNA-specific RT primer per the manufacturer’s instructions. PCR was performed for each RT sample using 1.33 uL of cDNA and Taqman™ Universal Master Mix II without Uracil-N-glycosylase (UNG). Controls included reaction mixtures without any reverse transcriptase, without template cDNA, and without master mix. PCR was performed in a 96-well standard (0.2mL) plate on a CFX96 Real-Time PCR Detection System (Bio-Rad Laboratories). Expression patterns were revealed using the delta-delta Ct method with SmsR1491 or SmsR847 as the normalization gene.

### Northern blotting

Total RNA was isolated as described above and diluted to 1 ug/uL. Samples containing 5 ug of RNA (5 uL) were combined with 5 uL of loading buffer (80% formamide, 10mM EDTA pH 8.0, 0.5mg/mL xylene cyanol, 0.5mg/mL bromophenol blue), incubated at 90°C for 4 minutes, and cooled on ice for 3 minutes. The 10 uL reaction mixture was loaded onto an 8% polyacrylamide/8M urea gel (National Diagnostics) alongside a Ambion Decades Markers System ladder that was radiolabeled per the manufacturer’s instructions (Thermo Fisher Scientific). Samples were resolved in 1X TBE at 300V for 1.25-1.5 h. RNA was transferred to a Zeta-Probe GT membrane (Bio-Rad) using a Hoefer TE Series transfer apparatus filled with 0.5X TBE buffer at 20 V overnight at 4℃. Membranes were UV cross-linked on both sides using a UV Crosslinker FB-UVXL-1000 (Thermo Fisher Scientific) (1200x100uJ/cm^2^) and pre-hybridized in hybridization tubes with 20 mL of ULTRAhyb-Oligo buffer (Thermo Fisher Scientific) at 45°C for 2 h with rotation. DNA probes complementary to the sequence of each sRNA were end-labeled with [*γ*-^32^P]dATP (Perkin-Elmer) and T4 polynucleotide kinase (New England BioLabs) as described before (28) (Haswell et al. 2013) with several alterations. First, 2 uL of a 20mM stock of the DNA target probe was combined with 13 uL nuclease free water and 2 uL T4 polynucleotide kinase buffer. The samples were boiled for 1 minute and cooled prior to adding 2 uL of [*γ*- ^32^P]dATP (Perkin-Elmer) and 1 uL T4 polynucleotide kinase. Following incubation, TE Select-D G-25 spin columns (Roche Applied Science) were used to remove unincorporated [*γ*-^32^P]dATP. Probes were heated at 90°C for 3 min and cooled on ice before adding to the hybridization buffer. Membranes were incubated overnight at 45°C with rotation and then washed twice with 2X SSC followed by three washes in 0.2X SSC with gentle agitation. The membranes were allowed to air dry before exposing them to Kodak BioMax film for up to 10 days at -80°C with an intensifying screen prior to autoradiography.

#### EMSA

Electromobility shift assays (EMSA) were performed as described previously (Haswell et al. 2013) to determine whether SloR binds to select sRNAs directly and whether the region of binding shares overlap with a proposed promoter region. Primers were designed to include the sRNA and its predicted promoter sequence, yielding a 201 bp amplicon for SmsR1532 and a 264 bp amplicon for SmsR1785 (Table 2). PCR amplification was performed with Q5 polymerase according to the manufacturer’s instructions (New England BioLabs) and the following thermal cycling conditions: initial denaturation at 98°C for 30s, followed by 35 cycles of denaturation at 98℃ for 10s, annealing at the optimal primer pair temperature for 30s, and extension at 72℃ for 30s, with a final extension time of 72℃ for 2 minutes. Amplicons were purified with the QIAquick PCR Purification Kit (Qiagen) according to the manufacturer’s instructions, confirmed by agarose gel electrophoresis, and quantified by NanoDrop spectrophotometry. Annealing of complementary oligonucleotides was performed by mixing 3.75 mM of the positive and negative DNA strands in annealing buffer in a total reaction volume of 60 uL and exposing the DNA mixture to the following cycling conditions in a thermal cycler: 95℃ for 2 min followed by 95℃ for 12 seconds for a total of 450 cycles with each cycle decreasing in temperature by 0.1℃.

The resulting amplicons and annealed oligos were end labeled with [*γ*-^32^P]dATP. Binding reactions were prepared as previously described (28) (Haswell et al. 2013) and incubated at room temperature for 20 minutes. EDTA was added to select reaction mixtures at a final concentration of 15 mM to chelate metal ions and determine if SloR binding is metal ion dependent. Samples were resolved on 12% non-denaturing polyacrylamide gels for 1,050-volt hours. Gels were exposed to Kodak BioMax film for up to 24 h at -80°C with an intensifying screen prior to autoradiography.

### sRNA target prediction

Both IntaRNA (83–86) and TargetRNA2 freeware (87) were used with default parameters to predict the downstream mRNA targets of *S. mutans* sRNAs at a genome-wide level (Table S13-16). KEGG Mapper (88, 89) was used to reveal the biological pathways in which the sRNAs might be involved. IntaRNA was also used to examine whether SmsR1532 binds to the *sloABCR* operon when each of the genes was entered manually; here, a seed region of at least 7 nt was used.

### Data availability

RNA-seq reads were deposited in NCBI BioProject under the accession number PRJNA976357.

## ACKNOWLEDGEMENTS

This research was supported by NIH grant DE014711-10 to G.A.S., by the Middlebury College Department of Biology, by the Middlebury Senior Research Project Supplement, and by the Vermont Biomedical Research Network’s Institutional Development Award (IDeA) from the National Institute of General Medical Sciences of the National Institutes of Health under grant number P20GM103449. The research reported in this publication is solely the responsibility of the authors and does not necessarily represent the official views of NIGMS or NIH. Next-generation sequencing was performed in the Vermont Integrative Genomics Resource Massively Parallel Sequencing Facility and was supported by the University of Vermont Cancer Center, Lake Champlain Cancer Research Organization, UVM College of Agriculture and Life Sciences, and the UVM Larner College of Medicine.

We acknowledge Gary Nelson for figure preparation, Dr. Robert Haney for perl scripts used in mining sRNA sequences in the literature, and Daniel Tetreault for assistance with the northern blots.

We have no conflicts of interest to declare.

## SUPPLEMENTARY MATERIAL

**Table S1.** Characterization of RNA-seq output. The number of small (18-50 nts) transcripts in each sRNA library ranged from 6 to 10 million reads, with the exception of 159_rep4 (>38 million). Most transcripts were of high quality and aligned with the *S. mutans* UA159 genome. Some mapped to IGRs and were selected for further investigation.

| Sample | Transcripts | Transcripts mapping to *S. mutans* genome | Transcripts mapping to *S. mutans* genome IGRs |
| --- | --- | --- | --- |
| UA159 replicate 1 | 6,991,074 | 6,459,185 | 1,238,805 |
| UA159 replicate 2 | 10,448,413 | 8,379,579 | 1,441,276 |
| UA159 replicate 3 | 10,760,504 | 10,275,360 | 2,142,103 |
| UA159 replicate 4 | 38,573,777 | 36,752,470 | 7,707,462 |
| GMS584 replicate 1 | 6,318,812 | 5,444,036 | 1,606,744 |
| GMS584 replicate 2 | 11,896,239 | 9,129,825 | 2,598,584 |
| GMS584 replicate 3 | 6,649,811 | 6,056,446 | 1,259,096 |
| GMS584 replicate 4 | 9,428,250 | 8,593,298 | 2,207,493 |
| Low Mn<sup>2+</sup> replicate 1 | 2,087,757 | 2,030,271 | 339,956 |
| Low Mn<sup>2+</sup> replicate 2 | 4,265,355 | 4,170,825 | 626,720 |
| Low Mn<sup>2+</sup> replicate 3 | 11,521,746 | 10,281,848 | 1,138,787 |
| High Mn<sup>2+</sup> replicate 1 | 9,766,429 | 9,546,823 | 1,689,262 |
| High Mn<sup>2+</sup> replicate | 8,938,525 | 8,749,294 | 1,436,625 |
| High Mn<sup>2+</sup> replicate | 12,879,486 | 11,628,161 | 1,468,840 |

**Table S2. sRNA_MASTER.fa.** Shown are the putative sRNA transcripts from streptococcal species and *Staphylococcus aureus* described in the literature. sRNA-seq results in the present study were compared to the published list of transcripts using NCBI BLASTN to determine whether any sRNAs predicted herein are conserved across species.

**Table S3.** Reports of *Streptococcus mutans* sRNAs in the literature. Shown is a compilation of all *S. mutans* sRNA reports in the published literature as of August 2022. Taken collectively, these reports describe a putative role for *S. mutans* sRNAs in growth, temperature and acid stress tolerance, carbohydrate utilization and metabolism, exopolysaccharide synthesis, adherence, biofilm formation, competence, and toxin-antitoxin systems. Importantly, sRNAs appear to be expressed across a variety of different clinical isolates of *S. mutans*.

| Article | Techniques | Conditions investigated | Key Findings | Number of sRNAs added to .fa file |
| --- | --- | --- | --- | --- |
| **Lee and Hong 2012 (90)**<br>Analysis of microRNA-size, small RNAs in *Streptococcus mutans* by deep sequencing | RNA-seq; validation with RT-PCR and northern blot; prediction of secondary structure with adjacent 60 nt mFold. | Transcripts 15-26 nt; UA159; grown in BHI; no information provided about timeframe, temperature, CO<sub>2</sub> levels, or growth-phase. | 922 unique transcripts that map to UA159 with secondary structure with the adjacent 60 nt; 36 transcripts with >100 reads; 6 verified via qPCR; msRNA-428 verified via northern blot. | 628 |
| **Xia et al. 2012 (91)**<br>Identification and Expression of Small Non-Coding RNA, L10-Leader, in Different Growth Phases of *Streptococcus mutans* | sRNA bioinformatic prediction with sRNA Predict, sRNASVM, and SIPHT; validation with qRT-PCR and northern blot; target prediction with sTarPicker and RNAPredator; analysis of putative target expression with qRT-PCR. | Transcripts 26 nt; UA159; growth phase; 2 strains with different adherence; pH 7.5 and 5.5; 0.1% and 1.0% glucose; 0.5% and 1.0% sucrose. | 40 sRNAs predicted by 2 or more methods. L10-leader sRNA validated and shown to be downregulated from log to stationary phase. Higher expression in the strain both with greater adherence and when under acid stress. No difference in various sucrose and glucose concentrations. The expression of 5 mRNA targets mirrored that of L10-Leader in growth phase and acid stress. | 40 |
| **Koyanagi and Levesque 2013 (41)**<br>Characterization of a *Streptococcus mutans* intergenic region containing a small toxic peptide and its cis-encoded antisense small RNA antitoxin | Validation with RT-PCR; RACE; northern blot; ChemiDoc XRS System and Quantity One software; toxicity assays; GFP-assay; persistence assay. | An unannotated Fst-like type 1 toxin-antitoxin system in UA159 IGR 176. | Confirmation of a type 1 toxin-antitoxin system, named Fst-Sm/srSm (~200 nt coding region and ~70 nt non-coding RNA). srSm binds to and represses Fst-Sm expression. The half-life of the *fst-Sm* mRNA is ~3x longer than that of the srSm. sRNA *fst-Sm* transcription confers toxicity in *E. coli* and *fst-Sm* overexpression confers toxicity in *S. mutans*, which can be neutralized by srSm expression. | 1 |
| **Zeng et al. 2013 (92)**<br>Gene regulation by CcpA and catabolite repression explored by RNA-seq in *Streptococcus mutans* | sRNA transcript prediction with RNAz, MUSCLE, Rho-independent terminator-based and transcriptional signal-based identifications; RNA-seq; target prediction with RNAplex and RNAplfold. | sRNAs in IGRs, length not provided; carbohydrate source (glucose or galactose), presence/absence catabolite control protein CcpA (UA159/TW1). | Differentially expressed transcripts matching predicted sRNAs: 5 UA159 glucose/galactose, 1 TW1 glucose/galactose; 2 glucose UA159/TW1; 2 galactose UA159/TW1. Interestingly, the 2 sRNAs galactose UA159/TW1 are regulated independently of surrounding genes. | sRNA sequences not provided |
| **Chylinski et al. 2013 (93)**<br>The tracrRNA and Cas9 families of type II CRISPR- Cas immunity systems | sRNA-seq; Cas9 sequence analysis with position-specific iterated (PSI) BLAST and BLASTClust; Cas9 multiple sequence alignment with MUSCLE, PSI-BLAST, and HHpred; Cas9 tree construction with FastTree. Analysis of CRISPR-Cas loci with CRSIPRdb database or CRISPRfinder tool, as well as BLASTP and/or KEGG database; *in silico* prediction of tracrRNA with Cector NTI software; promoter prediction with BDGP Neural Network Promoter Prediction program; tracrRNA Rho-independent terminator prediction with TransTermHP; tracrRNA multiple sequence alignment with MUSCLE; tracrRNA secondary structure RNAalifold. | Cas9, CRISPR-Cas, crRNA, tracrRNAs in various bacteria, including *S. mutans* UA159. | *S. mutans* utilizes a type II-A CRISPR-Cas9 system, meaning it requires tracrRNAs and a fourth cas gene which is *cas2*-like. TracrRNAs are a unique family of sRNAs required in type II CRISPR-Cas systems. Similar to many sRNAs, the tracrRNA family has conserved function despite not having conserved sequence. | 5 |
| **Liu et al. 2015 (94)**<br>The *Streptococcus mutans* irvA gene encodes a trans-acting riboregulatory mRNA | irvA deletion strain construction; DDAG assay; 5' RACE; mRNA stability assays; Western blots; *in vivo* AMT cross-linking; affinity purification of gbpC-irvA mRNA coprecipitates using 5'- biotinylated oligonucleotides; 5' "RACE walk"; *in vivo* and *in vitro* RNA EMSA; *in vitro* RNase J2 digestion of gbpC; FLAsH labeling and fluorescence microscopy; natural competence assay. | presence/absence of irvA. | The 5' UTR of irvA mRNA, which codes for a stress induced protein, is a trans-acting riboregulator. It binds to the gbpC mRNA (a critical activator of DDAG response) via a seed region, preventing degradation of the transcript by an RNase J2-mediated pathway. The IrvA protein functions independently to regulate natural competence. | 1 |
| **Liu et al. 2016 (95)**<br>Analysis of small RNAs in *Streptococcus mutans* under acid stress—a new insight for caries research | RNA-seq; target prediction with RNAhybrid; validation and investigation of sRNA and predicted mRNA target expression with qRT-PCR; assessment of bacterial growth characteristics and vitality assessment via absorbance at 600 nm and quantifying CFUs/volume. | Transcripts 18-50 nt; UA159; pH 7.5, 6.5, 5.5, and 4.5 at 0.5 hr, 1 hr, and 2 hrs. | 1879 sRNAs with at least 100 mean reads differentially expressed in acidic conditions. Investigation of 10 sRNAs and their predicted targets revealed correlation between sRNA expression and some mRNA targets at various pHs and times. | 1879 |
| **Mao et al. 2016 (96)**<br>The *rnc* gene promotes exopolysaccharide synthesis and represses the *vicRKX* gene expressions via microRNA-size small RNAs in *Streptococcus mutans* | Deletion and overexpression mutant, EPS, and biofilm assays; FGENESB and BPROM to predict operons and promoters; animal study examining cariogenicity of *rnc* mutants; measurement of *vicRKX* expression in *rnc* (+) and (-) strains with qRT-PCR; RNA-seq; target prediction with IntaRNA; validation with validation step-loop qRT-PCR. | IGR and antisense transcripts 18-150 nt; presence/absence of *rnc* (UA159/Smurnc). | *rnc* in *S. mutans* significantly positively regulates EPS synthesis and alters biofilm formation; cariogenicity decreased in *rnc*- strain; *rnc* represses *vicRKX*, apparently using 3 msRNAs at post-transcriptional level. | 3 |
| **Liu et al. 2017 (97)**<br>Analysis of sucrose-induced small RNAs in *Streptococcus mutans* in the presence of different sucrose concentrations | RNA-seq; target prediction with RNAhybrid; validation and investigation of sRNA and predicted mRNA target expression in different sucrose concentrations with qRT-PCR; bacterial vitality assessment by quantifying CFUs/volume; adherence analysis; polysaccharide synthesis analysis. | Transcripts 18-50 nt. UA159; 1% and 5% sucrose medium. | 22 differentially expressed sRNA candidates with >100 avg reads. 2 upregulated, 20 downregulated with respect to 1% sucrose. The predicted target gene for one particular sRNA is upregulated in 1% sucrose. Synthesis of water-insoluble glucans significantly higher in 5% sucrose. | 2125 |
| **Zhu et al. 2017 (98)**<br>Characterization of acid-tolerance-associated small RNAs in clinical isolates of *Streptococcus mutans*: Potential biomarkers for caries prevention | Use of Liu et al. 2016 RNA-seq data; prediction of target mRNAs and their pathways with RNA predictor and KEGG; assessment of acid tolerance with acid killing assay; assessment of sRNA133474 and predicted target mRNA with qRT-PCR. | One specific sRNA, sRNA133474; UA159; pH 5.5 and 7.5; 3 different growth phases; 20 clinical isolates. | Of 6 sRNAs tested in UA159 and 3 clinical strains, 3 differentially expressed. sRNA133474 most highly downregulated in pH 5.5. Found to be sig. downregulated at pH 5.5 in 10 clinical isolates as well. In these 10 strains, 2 different patterns: in 6 strains sRNA133474 was downregulated in early log to early stationary phase, in 4 strains, sRNA133474 downregulated until early stationary phase, when it was upregulated. Expression of 3 predicted target mRNAs (all within TCS pathway) were (-) correlated with expression of sRNA133474. In short, *S. mutans* may use sRNA133474 as an intermediary in TCS pathway to aid in acid tolerance. | Liu et al. 2016 data |
| **Zhu et al. 2018 (99)**<br>High-throughput sequencing identification and characterization of potentially adhesion-related small RNAs in *Streptococcus mutans* | RNA-seq; target mRNA prediction with IntaRNA; pathway prediction with GO and DAVID; validation and investigation of sRNA and predicted mRNA target expression with qRT-PCR; adherence analysis; biofilm assessment with fluorescence staining. | Transcripts 18-150 nt; 1% and 0% sucrose; UA159 and 100 clinical strains; assessment of adhesion in 4 different adhesion conditions (0 % sucrose, 0.5 % sucrose, 1% sucrose and 2% sucrose) for 1, 2, 3 or 4h. | There were 736 differentially expressed sRNAs, 352 antisense, 384 in IGRs. Top 7 validated via PCR in UA59, 2 in 100 clinical isolates, and 2 conserved across *Streptococci*. (+) correlation between sRNAs and adhesion of 100 clinical strains. | Unable to convert to excel file |
| **Liu et al. 2019a (100)**<br>Effect of Different Glucose Concentrations on Small RNA Levels and Adherence of *Streptococcus mutans* | RNA-seq; target prediction analysis with KEGG; validation and investigation of sRNA and predicted mRNA target expression in different sucrose concentrations with qRT-PCR; pH drop was assessed in 1% and 5% glucose, as was adherence capacity and polysaccharide synthesis (including WIGs and IPSs). | Transcripts 18-50 nt; UA159; 1% and 5% glucose | pH drop and adherence capacity in 1% glucose were similar to those in 5% sucrose in Liu et al. 2016. Adherence capacity higher in 1% glucose. There were 2149 putative sRNAs with >100 avg reads, ~½ map to acid-induced sRNAs (Liu et al. 2016), >½ map to sucrose-induced sRNAs (Liu et al. 2017), 11 map to msRNAs in Lee and Hong 2012). There were 581 differentially expressed, 268 upregulated, 313 downregulated in 1% glucose; 43 verified, 22 sig. upregulated, 17 sig. downregulated. Predicted mRNA target genes did not show differential expression. |  |
| **Liu et al. 2019b (101)**<br>Insight into the Effect of Small RNA smn225147 on Mutacin IV in *Streptococcus mutans* | Use of Liu et al, 2019a RNA-seq data; target mRNA prediction with RNAhybrid and RNAPredator; determination of KEGG pathways associated with smn225147 with DAVID and antiSMASH bacterial version; selection of clinical isolates with high and no mutacin IV production with bacteriocin assay; assessment of expression of smn225147 and predicted target *comD* in high and low mutacin IV strains with qRT-PCR; assessment of smn225147 impact on *comD* expression the addition of mimic smn225147 via electroporation and qRT-PCR. | Clinical isolates (20) with high and no production of mutacin IV, a bacteriocin. | Levels of ComD, a mutacin IV-formation associated gene, significantly higher in high-mutacin IV strains, smn225147 higher in no-mutacin IV strains. At 400-fold overexpression of smn225147, ComD levels were decreased, but at 1400-fold overexpression of smn225147, ComD levels were increased. smn225147 appears to have a two-way regulatory effect on *comD* expression, although no result was sig. Appears to be a 2-way regulatory effect of smn225147 on *comD*, but weak evidence of effect on mutacin-IV. | 50 |
| **Yin et al. 2020 (102)**<br>Small noncoding RNA sRNA0426 is involved in regulating biofilm formation in *Streptococcus mutans* | Use of Zhu et al. 2018 RNA-seq data; assessment of sRNA expression in planktonic/sessile states with qRT-PCR to evaluation of biofilm biomass with crystal violet assay; EPS production analysis in UA159 and 10 clinical strains with confocal lens scanning microscopy; prediction of target mRNAs with RNAfold. | Transcripts 18-50 nt; sRNAs in biofilm formation, specifically expression in planktonic versus sessile states. | Of the 20 sRNAs tested, 18 detected, 14 were upregulated and 4 downregulated with respect to planktonic status. One transcript, sRNA0426, had a strong (+) relationship with biofilm formation and had a (+) correlation with EPS production. Predicted targets were particularly involved in carbon metabolism; 3 targets involved in EPS synthesis (+) correlated with sRNA0426, 1 (-) correlated. | Zhu et al. 2018 data, i.e., not available |
| **Krieger et al. 2022 (37)**<br>Genome-wide identification of novel sRNAs in *Streptococcus mutans* | Growth assays; RNA-seq; prediction of secondary structure with RNAalifold; ARNold and TranTermHP for terminator prediction; assessment of expression under 4 conditions of stress via qRT-PCR; *in vitro* transcription; northern blotting to confirm transcription; EMSA with mutagenesis to confirm sRNA-mRNA binding; gene deletion and complementation; 5' and 3' RACE; covariance modelling; target prediction with TargetRNA2 and IntaRNA | Presence of sugar-phosphate (10%), hydrogen peroxide exposure, high temperature, and low pH stress; SmsR4 existence in other streptococcal species; presence/absence SmsR4 in sorbitol-containing media | 15 novel sRNAs were uncovered, some of which appear to be homologs of sRNAs in other bacteria. Some sRNAs contained predicted ORFs, indicative of them being dual-function sRNAs. The 15 sRNAs respond to various environmental stressors (presence of sugar-phosphate, hydrogen peroxide exposure, high temperature, low pH). SmsR4, which does exist in other streptococcal species although perhaps in different a capacity, aids in cell growth in the presence of sorbitol apparently through regulation the enzyme IIA (EIIA) component of the sorbitol phosphotransferase system to which is binds. | 15 |

**Table S5.** sRNAs subject to SloR and/or manganese control. All sRNAs that demonstrated at least a 2-fold change in expression are listed in order of most positive (upregulated in UA159 compared to GMS584 or upregulated in low manganese compared to high manganese) to most negative (upregulated in GMS584 compared to UA159 or upregulated in high manganese compared to low manganese) per the RNA-seq output.

| Name<sup>a</sup> | Length (nt)<sup>b</sup> | Strand | 5' coordinate<sup>b</sup> | 3' coordinate<sup>b</sup> | 5' adjacent gene | 3' adjacent gene | Expression UA159/GMS584<sup>c</sup> | Expression high/low Mn<sup>d</sup> | Confirmation<sup>e</sup> | Previous prediction<sup>f</sup> |
| --- | --- | --- | --- | --- | --- | --- | --- | --- | --- | --- |
| SmsR929 | 25 | + | 1335086 | 1335110 | SMU_1405c (cas9) | SMU_1406c (nfr1_3, NADH-dependent flavin reductase subunit 1) | ↑ | P, NDE | NB, TM | srRNA30 (91) |
| SmsR344 | 21 | + | 1793797 | 1793817 | SMU_1912c (hypothetical protein) | SMU_1913c (putative immunity protein, BLpL-like) | ↑ | NP |  | srn821610 (95, 97) |
| SmsR284 | 20 | + | 1793786 | 1793805 | SMU_1912c (hypothetical protein) | SMU_1913c (putative immunity protein, BLpL-like) | ↑ | P, NDE |  |  |
| SmsR370 | 23 | + | 1793706 | 1793728 | SMU_1912c (hypothetical protein) | SMU_1913c (putative immunity protein, BLpL-like) | ↑ | P, NDE |  |  |
| <u>SmsR1785/SmsR3452*</u> | 66 | + | 211469 | 211534 | SMU_219 (hypothetical protein) | SMU_220c (hypothetical protein) | ↑ | P, NDE | NB, TM | srSm (41) |
| SmsR424 | 18 | - | 1793871 | 1793854 | SMU_1913c (putative immunity protein, BLpL-like) | SMU_1912c (hypothetical protein) | ↑ | NP |  |  |
| SmsR1287 | 18 | + | 1794409 | 1794426 | SMU_1913c (putative immunity protein, BLpL-like) | SMU_1914c (hypothetical protein) | ↑ | P, NDE |  |  |
| SmsR1071 | 18 | - | 1794424 | 1794407 | SMU_1914c (hypothetical protein) | SMU_1913c (putative immunity protein, BLpL-like) | ↑ | P, NDE |  |  |
| SmsR306 | 18 | - | 1758111 | 1758094 | mutY (putative A/G-specific DNA glycosylase) | SMU_1862 | ↑ | P, NDE |  |  |

| SmsR917 | 33 | - | 1864393 | 1864361 | pbp1b (SMU_1991) | rpoB (SMU_1990) | ↑ | P, NDE |  |  |
| --- | --- | --- | --- | --- | --- | --- | --- | --- | --- | --- |
| SmsR563 | 18 | + | 1254882 | 1254899 | SMU_1332c (putative transposase) | sfp (putative phosphopantethe inyl transferase) | ↑ | P, NDE |  |  |
| SmsR1491 | 26 | + | 520383 | 520408 | divIVA (putative cell division protein DivIVA) | SMU_558 (isoleucine-tRNA synthetase) | ↓ | P, NDE | NB, TM |  |
| SmsR1347 | 37 | + | 44769 | 44805 | SMU_41 (hypothetical protein) | SMU_42 (conserved hypothetical protein) | ↓ | P, NDE |  | srn19368 (95) |
| SmsR1718 | 25 | + | 935481 | 935505 | SMU_988 (putative cardiolipin synthase) | asd (aspartate-semialdehyde dehydrogenase) | ↓ | P, NDE |  | srn447908 (95, 97) |
| SmsR1656 | 18 | + | 1224110 | 1224127 | SMU_1297 (conserved hypothetical protein) | r131 (50S ribosomal protein L31) | ↓ | P, NDE |  |  |
| SmsR1480 | 23 | - | 1758402 | 1758380 | mutY (putative A/G-specific DNA glycosylase) | SMU_1862 (hypothetical protein) | ↓ | P, NDE |  |  |
| SmsR850/SmsR7533 | 25 | - | 1190856 | 1190832 | SMU_1258c (possible restriction endonuclease) | SMU_1257c (conserved hypothetical protein) | ↓ | ↓ | NB |  |
| SmsR1532* | 30 | + | 183345 | 183373 | SloR | SMU_187c | ↓ | P, NDE | NB, TM |  |
| SmsR1765 | 43 | + | 497896 | 497938 | SMU_530c (conserved hypothetical protein) | SMU_531 (putative corismate mutase) | ↓ | P, NDE |  | SmsR6 (37) |
| Name<sup>a</sup> | Length<sup>b</sup> (nt) | Strand | 5' coordinate<sup>b</sup> | 3' coordinate<sup>b</sup> | 5' adjacent gene | 3' adjacent gene | Expres-<br>sion UA159/<br>GMS584<sup>c</sup> | Expres-<br>sion high/<br>low Mn<sup>2+</sup><sup>c</sup> | Confirm<br>ation<sup>d</sup> | Previous<br>prediction<sup>e</sup> |
| SmsR7533/SmsR850 | 25 | - | 1190856 | 1190833 | SMU_1258c (possible restriction endonuclease) | SMU_1257c (conserved hypothetical protein) | ↓ | ↓ | NB |  |
| SmsR3509 | 34 | - | 995514 | 995481 | SMU_1047c CDS (hypothetical protein) | SMU_1046c CDS (putative GTP pyrophosphokinase) | P, NDE | ↑ |  | srn471693/<br>srn471694 (95); SmsR7 (37) |
| SmsR862 | 36 | - | 202844 | 202809 | SMU_206c (hypothetical protein) | SMU_205c (hypothetical protein) | P, NDE | ↑ |  | srn90412 (95) |
| SmsR18621 | 18 | - | 441409 | 441392 | SMU_473 CDS (conserved hypothetical protein) | SMU_472 CDS (conserved hypothetical protein; possible N6-adenine-specific DNA methylase) | P, NDE | ↑ |  |  |
| SmsR16813 | 27 | - | 995472 | 995446 | SMU_1047c | SMU_1046c | P, NDE | ↑ |  |  |

|  |  |  |  |  | CDS<br>(hypothetical<br>protein) | CDS (putative<br>GTP<br>pyrophosphokina<br>se) |  |  |  |  |
| --- | --- | --- | --- | --- | --- | --- | --- | --- | --- | --- |
| SmsR11680 | 24 | + | 303839 | 303862 | SMU_318 CDS<br>(putative<br>hippurate<br>hydrolase) | SMU_320 CDS<br>(putative 5-formyltetrahydro<br>folate cyclo-<br>ligase) | P, NDE | ↑ |  |  |
| SmsR17090 | 42 | - | 1161933 | 1161892 | SMU_1220c<br>CDS (conserved<br>hypothetical<br>protein) | SMU_1219c<br>CDS (conserved<br>hypothetical<br>protein) | P, NDE | ↑ |  |  |
| SmsR19262 | 36 | - | 1327935 | 1327900 | SMU_1399 CDS<br>(hypothetical<br>protein) | SMU_1398 CDS<br>(putative<br>transcriptional<br>regulator) | P, NDE* | ↑ |  | srn628416-22<br>(95) |
| SmsR22851 | 42 | - | 892757 | 892716 | mvaA CDS<br>(putative<br>hydroxymethylgl<br>utaryl-CoA<br>reductase) | SMU_941c CDS<br>(conserved<br>hypothetical<br>protein) | NP | ↑ |  |  |
| **<u>SmsR3452/S</u>**<br>**<u>msR1785\*</u>** | 66 | + | 211469 | 211534 | SMU_219<br>(hypothetical<br>protein) | SMU_220c<br>(hypothetical<br>protein) | ↑ | P, NDE | NB, TM | srSm (41) |

<sup>a</sup>Small RNAs were named SmsR for *S. mutans* small RNA. Numbers correspond to the read number assigned in RNA-seq data processing. sRNAs present in both the UA159/GMS584 and low/high Mn libraries appear in **boldface type**. sRNAs that are differentially expressed in both sets of libraries are in **boldface type and are underlined**. SmsR1491 and SmsR847 are included in this table and italicized; since they were not differentially expressed, they were used as endogenous controls. The stars indicate sRNAs that were selected for further analysis.
<sup>b</sup>The 5' and 3' ends of sRNAs were identified using visual scans of RNA-seq alignment files for coordinates where dramatic changes in the number of reads that mapped per nucleotide were visualized.
<sup>c</sup>sRNA expression was analyzed relative to expression in UA159 (i.e., sRNAs denoted as upregulated indicate higher expression in UA159, while sRNAs denoted as downregulated indicate higher expression in GMS584). P, NDE indicates that the sRNA is present but not differentially expressed in the SloR libraries, while NA indicates that the sRNA is not present in the SloR libraries.
<sup>d</sup>sRNA expression was analyzed relative to expression in conditions of low manganese (i.e., sRNAs denoted as upregulated indicate higher expression in low manganese, while sRNAs denoted as downregulated indicate higher expression in high manganese). P, NDE indicates that the sRNA is present but not differentially expressed in the manganese libraries, while NP indicates that the sRNA is not present in the manganese libraries.
<sup>e</sup>Confirmation of sRNAs in this or another study by northern blot (NB) or Taqman small RNA assay (TM).
<sup>f</sup>Existing and previously predicted sRNAs that overlap our results.

**Table S6. Complete representative Streptococcus genomes used in the creation of a phylogenetic tree.**

**Table S7. Complete Streptococcus genomes used in BLASTN analysis and covariance modeling.**

**Table S8. Query sequences used in BLASTN analysis and covariance modeling.**

**Table S9. SloR BLASTN results.**

**Table S10. SmsR1532 BLASTN results.**

**Table S11. SmsR1785 covariance modeling results.**

**Table S12. Summary of SloR and SmsR1532 BLASTN and SmsR1785 covariance modeling results.**

**Table S13. Top 100 target genes for SmsR1532 predicted by IntaRNA.**

**Table S14. All target genes for SmsR1532 predicted by TargetRNA2.**

**Table S15. All target genes for SmsR1785 predicted by IntaRNA.**

**Table S16. All targets for SmsR1785 predicted by TargetRNA2.**

**Figure S1.**
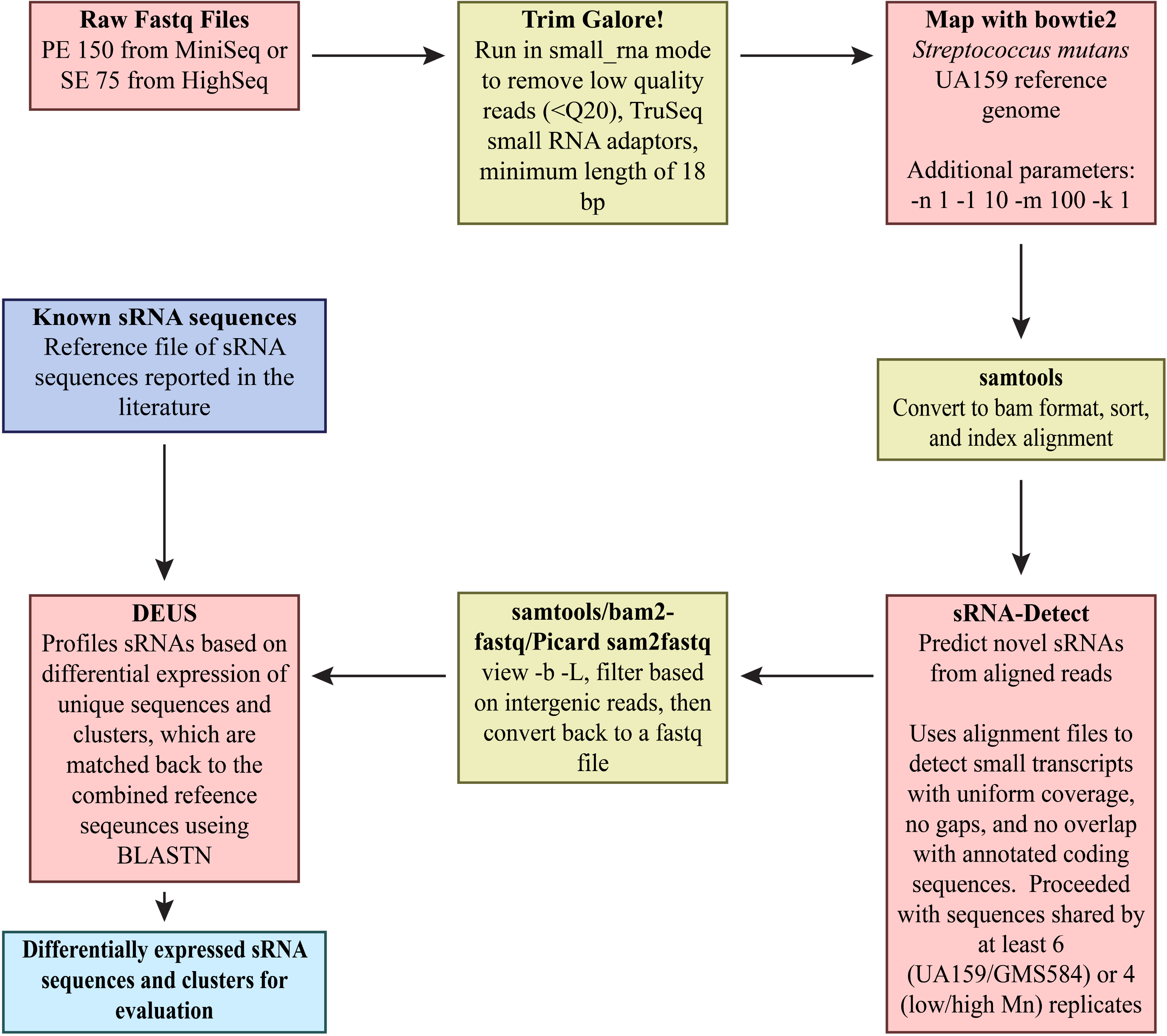
Workflow used for processing the RNA-seq output.

**Figure S2.**
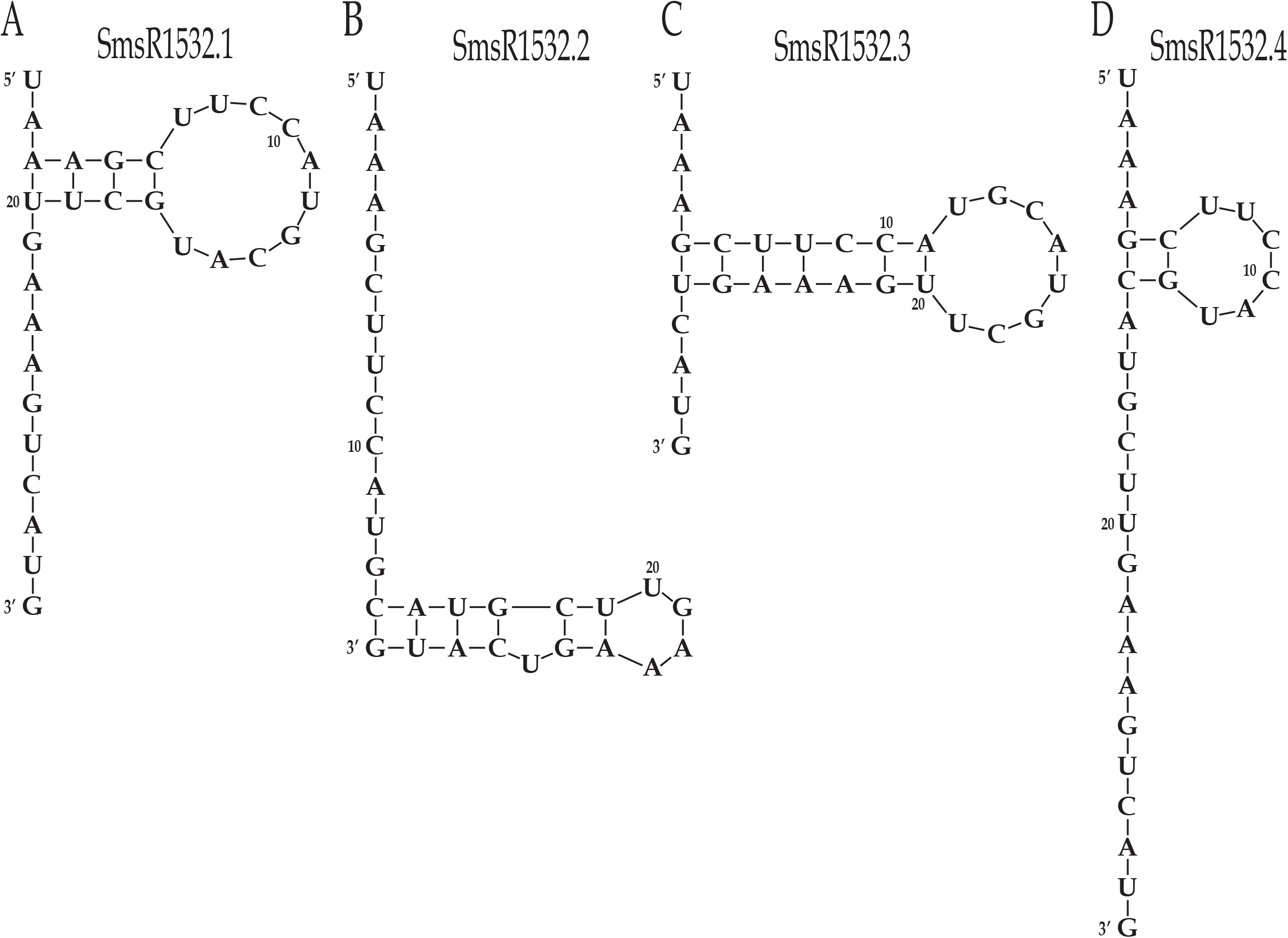
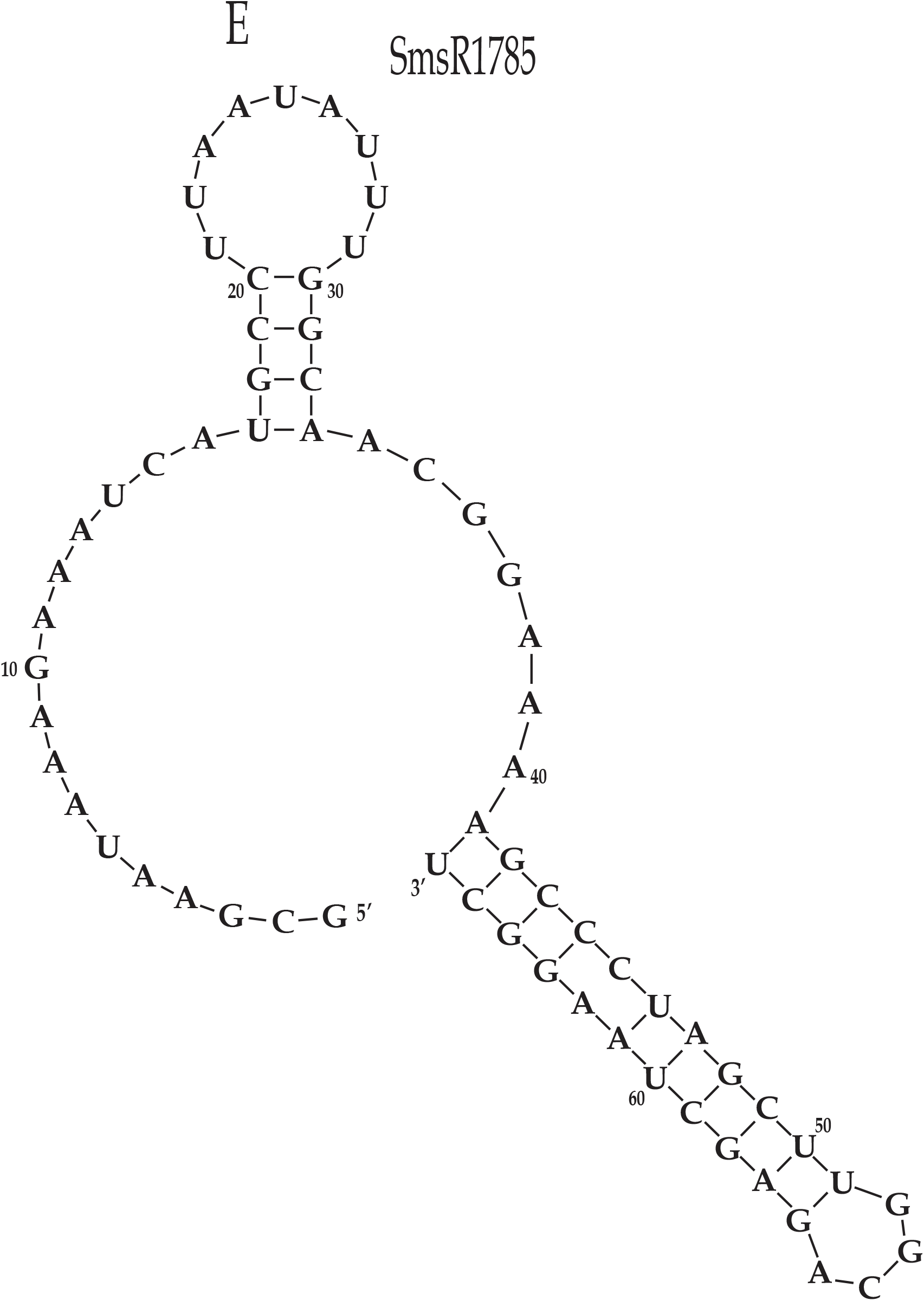
Predicted secondary structures for SmsR1532 and SmsR1785. mFold V3.6 was used to predict the best fit secondary structures for both SmsR1532 and SmsR1785. mFold predicted 4 secondary structures for SmsR1532 (SmsR1532.1, 1532.2, 1532.3 and 1532.4) and 1 structure for SmsR1785. All structures have a stable stem-loop structure which is characteristic of sRNAs. The lower the ΔG, the more likely the specific folding pattern is to occur. **A)** Shown is the predicted best fit secondary structure for SmsR1532 (30 nt) that has a stable stem-loop and a ΔG of -2.40 kcal/mol. **B-D)** These SmsR1532 predicted secondary structures (30 nt) all have stable stem loops and share a ΔG value of -1.90 kcal/mol. **E)** The predicted secondary structure for SmsR1785 (66 nt) forms a stable loop and has a ΔG of -17.90 kcal/mol.

**Figure S3.**
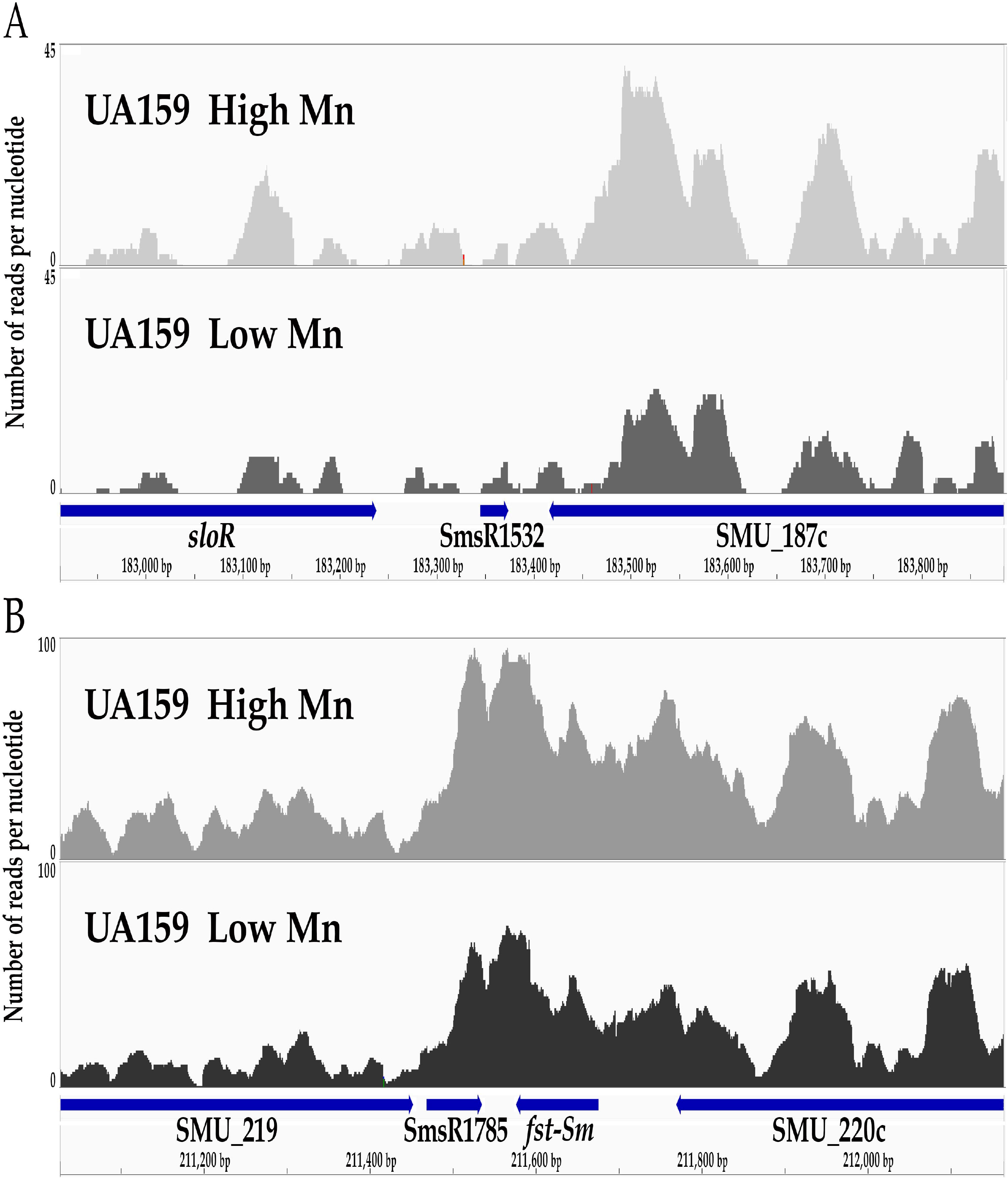
Coverage plots for *S. mutans* SmR1532 and SmsR1785 sRNAs in conditions of low or high manganese. sRNA-seq reads in the *S. mutans* UA159 wild-type strain grown in a medium adjusted with MnSO4 to low (1.7 uM) or high (75 uM) concentrations are shown. The y-axes indicate the number of RNA-seq reads that mapped to each nucleotide. The genome, gene, and sRNA annotations are provided below the coverage plots. **A)** SmsR1532 is transcribed as a 30 nt transcript from the IGR downstream of the *sloR* gene. **B)** SmsR1785 is trascribed as a 66 nt sRNA from the IGR downstream of the *fst-Sm* toxin gene. This sRNA has been previously described as an srSm antitoxin sRNA by Koyanagi and Levesque (41).

**Figure S4.**
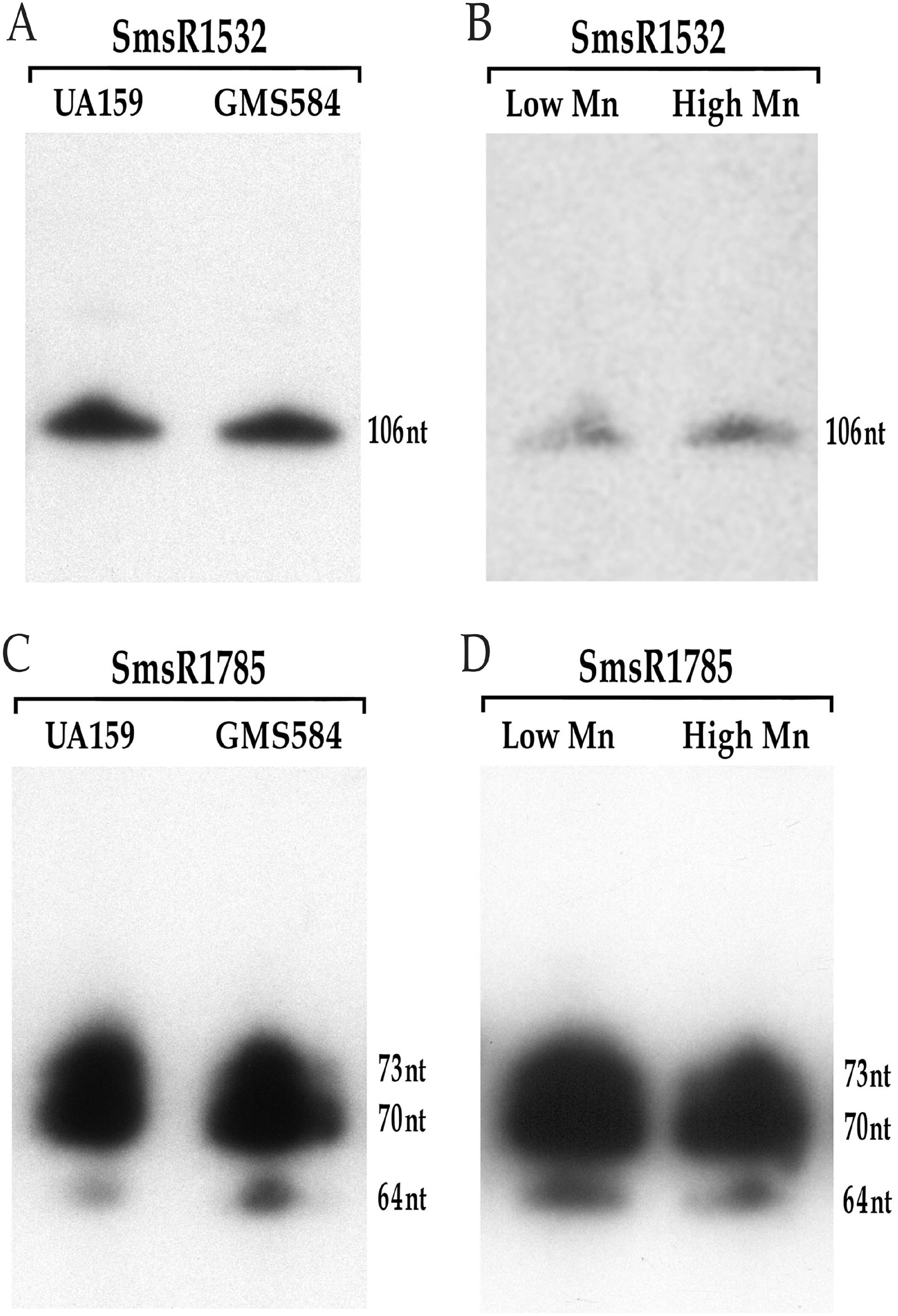
SmsR1532 and SmsR1785 derive from the processing of large transcripts. Northern hybridization experiments were performed with total RNA isolated from *S. mutans* UA159 and GMS584 cultures grown in BHI medium, and from UA159 grown in SDM containing low versus high MnSO4 concentrations. Radiolabeled single-stranded DNA oligonucleotides complementary to the sRNA sequence (Table 2) were used as probes to elucidate the size of the SmsR1532 and SmsR1785 transcripts. Transcripts were observed for *S. mutans* UA159 and GMS584 grown in BHI, and for UA159 grown in a semi-defined medium supplemented with high versus low manganese. **A and B)** The SmsR1532 probe hybridized with a 106 nt transcript in UA159 and GMS584, and in UA159 grown in high versus low manganese. This suggests RNA processing is necessary to generate the 30 nt sRNA that was revealed by RNA-seq. **C and D)** The SmsR1785 probe hybridized to 73 nt, 70 nt and 66 nt transcripts in UA159 and GMS584, and in UA159 grown in high versus low manganese. This supports processing of the 73 nt and 70 nt precursor transcripts to generate the 66 nt sRNA that was revealed by RNA seq.

